# SWR1 is recruited to activated ABA response genes to maintain gene body H2A.Z in *Arabidopsis thaliana*

**DOI:** 10.1101/2024.07.14.603444

**Authors:** Ellen G. Krall, Roger B. Deal

## Abstract

The histone variant H2A.Z is important for transcriptional regulation across eukaryotes, where it can alternately promote or repress transcription. In plants, actively transcribed genes show H2A.Z enrichment in nucleosomes immediately downstream of the transcription start site (TSS), while silent genes show H2A.Z enrichment across the gene body. Previous work showed that silent genes responsive to temperature and far-red light lose gene body H2A.Z upon activation, but whether H2A.Z loss is generally required for transcription is not clear. We profiled H2A.Z and components of its deposition complex, SWR1, before and after treating *Arabidopsis thaliana* with the hormone abscisic acid (ABA). Our results show that transcribed genes with TSS-enriched H2A.Z have high SWR1 binding at steady-state, indicating continuous replacement of H2A.Z, while silent genes with gene body H2A.Z show lower SWR1 binding. Surprisingly, upon ABA treatment, thousands of previously silent genes activate, coincident with recruitment of SWR1 and retention of gene body H2A.Z enrichment. We also found that the SWR1-interacting protein MBD9 is not required for SWR1 recruitment to activated genes. These results provide new insights into the relationship between H2A.Z and transcription and the mechanics of H2A.Z targeting to chromatin.

## Introduction

In the eukaryotic nucleus, histone proteins drive DNA organization by binding and condensing DNA into chromatin. The basic unit of chromatin is the nucleosome, where ∼147 base pairs of DNA are wrapped around a histone protein octamer composed of one H3/H4 histone tetramer flanked by two H2A/H2B histone dimers. Histones function to condense DNA, but their occupancy is also a physical barrier to DNA access. This occupancy prevents trans-acting factors from binding to regulatory elements depending on a cell’s developmental, environmental, and temporal state.

The modulation of DNA-histone binding is therefore crucial to promote proper transcription and other DNA-templated activities at any given moment within the eukaryotic cell. Chromatin remodeling complexes directly modify DNA-histone interactions by hydrolyzing ATP to slide, move, and eject nucleosomes as well as exchange canonical histones for specialized histone variants^1^. H2A.Z is a well-conserved histone variant of histone H2A deposited by the chromatin remodeler SWI2/SNF2-Related 1 (SWR1) ^2–4^.

The consequences of H2A.Z loss or misregulation on an organism are severe. H2A.Z misregulation contributes to the loss of cellular identity and the proliferation of cancer cells in melanoma, breast cancer, pancreatic ductal adenocarcinoma, prostate cancer, and hepatocellular carcinoma ^5–10^. Its loss is also embryonic lethal in many organisms including *C. elegans, Drosophila melanogaster,* and mammals ^11–15^. Plants, including the model plant *Arabidopsis thaliana* (Arabidopsis), tolerate the absence of H2A.Z but show pleiotropic phenotypes including early flowering, abnormal leaf development, reduced fertility, and an inability to respond appropriately to abiotic and biotic stressors ^16–18^. This makes Arabidopsis an ideal model organism to study the consequences of H2A.Z loss beyond embryonic development.

The effects of H2A.Z misregulation stem from its dual role in both positively and negatively regulating transcription, which are related to its deposition relative to occupied genes ^4,11,19^. When H2A.Z is enriched in nucleosomes flanking the Transcription Start Site (TSS), it creates conditions favorable for transcription. The first nucleosome downstream of the TSS (+1 nucleosome) is an energetic barrier to RNA polymerase, and incorporation of H2A.Z can lower this barrier by creating more mobile, easily unwrapped nucleosomes ^20,21^. When H2A.Z enrichment skews to cover whole gene bodies, it has a repressive effect ^16,22^. Genes that have more gene body H2A.Z are more responsive to environmental and developmental cues in plants, human cell lines, embryonic stem cells, CD4^+^ T lymphocyte cells, and intestinal epithelial cells ^22–27^. Additionally, these responsive genes are targets of Polycomb Repressive Complexes 1 and 2 (PRC1 and 2), which further mark these genes with post-translational histone modifications (PTMs) H2AK121ub and H3K27me3, respectively, establishing a repressed but responsive chromatin environment ^28,29^.

Post-translational modifications of H2A.Z itself further distinguish the two observed profiles of H2A.Z. TSS-proximal H2A.Z is often acetylated at N-terminal lysines K4, K7 and K11 ^30,31^. Acetylation of H2A.Z has been shown to contribute to gene activation in many contexts across eukaryotes and during tumorigenesis ^32–35^. Gene body associated H2A.Z (gbH2A.Z), on the other hand, is often monoubiquitinated at its C-terminal end at multiple lysines ^29–31^. This modification, as with canonical histone modification H2AK121ub, is deposited by PRC1 and contributes to gene repression ^29,36^. Additionally, gbH2A.Z is generally associated with the PRC2 mark, H3K27me3, where this PTM appears to be dependent on the presence of H2A.Z ^37^.

It is reasonable to assume that these two distinct profiles of H2A.Z enrichment, TSS-proximal H2A.Z and gbH2A.Z, are interconnected. Upon gene activation, gbH2A.Z may be lost as genes shift to having only TSS-proximal H2A.Z. Once repression is re-established, gbH2A.Z would also then be re-established. This notion is supported by experiments showing that H2A.Z is lost from genes responsive to far-red light, heat stress, and ethylene in plants once activated ^38–40^. However, it remains unclear if the distribution of H2A.Z on a gene is a feature that drives transcriptional regulation or simply reflects the present transcriptional status of the gene.

The establishment and maintenance of these two distinct H2A.Z profiles must depend at least in part on the H2A.Z deposition complex, SWR1. However, how SWR1 is targeted specifically to certain genes to deposit H2A.Z is unclear. SWR1 is a multi-subunit chromatin remodeler first identified in *S. cerevisiae*, and many of the 13 core subunits are conserved across eukaryotes ^4^. The ATP-hydrolyzing catalytic subunit is known as Domino in Drosophila, SRCAP/TIP60 in vertebrates, and PHOTOPERIOD-INDEPENDENT EARLY FLOWERING 1 (PIE1) in plants ^41^. Previous work has established that SWR1 is recruited to nucleosome-depleted regions (NDR) in organisms including yeast and mouse embryonic stem cells ^42–44^. Core components of SWR1 lack DNA-binding specificity, and therefore SWR1 interacts transiently with a variety of other factors that can directly affect its localization ^41,45^. However, the general and specific mechanisms governing the targeting of SWR1 by various factors remain unclear.

There is evidence that Bromodomain-containing accessory subunits are specifically involved in TSS-associated H2A.Z patterning on genes. For example, the yeast BromoDomain Factor 1 (BDF1) may bind to hyperacetylated histones around the TSS of active genes to recruit SWR1 ^46^. Similarly in Arabidopsis, the plant-specific METHYL-CPG-BINDING DOMAIN-CONTAINING PROTEIN 9 (MBD9) protein was proposed to bind to acetylation marks to recruit SWR1 for H2A.Z deposition ^47,48^. Our group as well as others demonstrated that MBD9 interacts with SWR1 and is necessary for proper H2A.Z accumulation in Arabidopsis chromatin ^48–50^. These publications demonstrated that plants lacking MBD9 had a reduction of H2A.Z enrichment at a subset of euchromatic sites compared to WT plants. However, it is still unclear if MBD9 is directly involved in SWR1 recruitment or if its effects on H2A.Z levels are indirect.

Here, we developed antibodies against the Arabidopsis SWR1 catalytic subunit, PIE1, and interactor MBD9 to better understand their localization in the Arabidopsis genome relative to H2A.Z. We also employed exogenous application of the plant hormone Abscisic Acid (ABA) to assess how their localizations change during a widespread transcriptional activation event. We further analyzed MBD9 deficient plants and induced an MBD9 Bromodomain-disrupting point mutation to define the role of MBD9 in SWR1 recruitment and subsequent H2A.Z deposition.

Our results indicate that silent genes with gbH2A.Z enrichment have much lower levels of the SWR1 ATPase PIE1 and interacting protein MBD9 under steady-state conditions compared to active genes with TSS-proximal H2A.Z enrichment. Upon ABA treatment, we see that PIE1 and MBD9 are recruited to previously silent ABA responsive genes with gbH2A.Z, but this surprisingly results in only a subtle decrease in H2A.Z enrichment on gene bodies. Parallel experiments in *mbd9-1* plants show that PIE1 is still recruited to newly activated genes in the absence of MBD9, indicating that MBD9 is not a targeting factor for SWR1 in this context. We also provide evidence that the MBD9 Bromodomain is dispensable to the role of MBD9 in the maintenance of H2A.Z levels. Taken together, this work suggests that SWR1 plays an active role in maintaining H2A.Z both at genes with TSS-enriched H2A.Z and normally silent genes with gene body H2A.Z enrichment. SWR1, along with interacting protein MBD9, is normally localized to active genes to continuously replace H2A.Z in the unstable TSS-proximal nucleosomes. Upon transcriptional activation of gbH2A.Z-enriched ABA responsive genes, SWR1 moves to these newly activated genes to maintain gene body H2A.Z, indicating that H2A.Z removal is not a prerequisite for activation of all gbH2A.Z-silenced genes. We also provide evidence that MBD9 and specifically the MBD9 Bromodomain, are not required for SWR1 targeting at these responsive genes.

## Results

### Genes with TSS-proximal H2A.Z have more PIE1 and MBD9 binding at steady state

To better understand the localization of SWR1 and MBD9 relative to H2A.Z, we utilized antibodies raised against unique peptide sequences of PIE1 and MBD9. These antibodies specifically recognize their respective proteins via Western blot (**Supp Fig. 1**), and to examine their localizations in the genome, we performed Chromatin Immunoprecipitation followed by Sequencing (ChIP-Seq) in 7-day-old WT Arabidopsis seedlings using these antibodies in addition to an antibody against H2A.Z.

**Figure 1.**
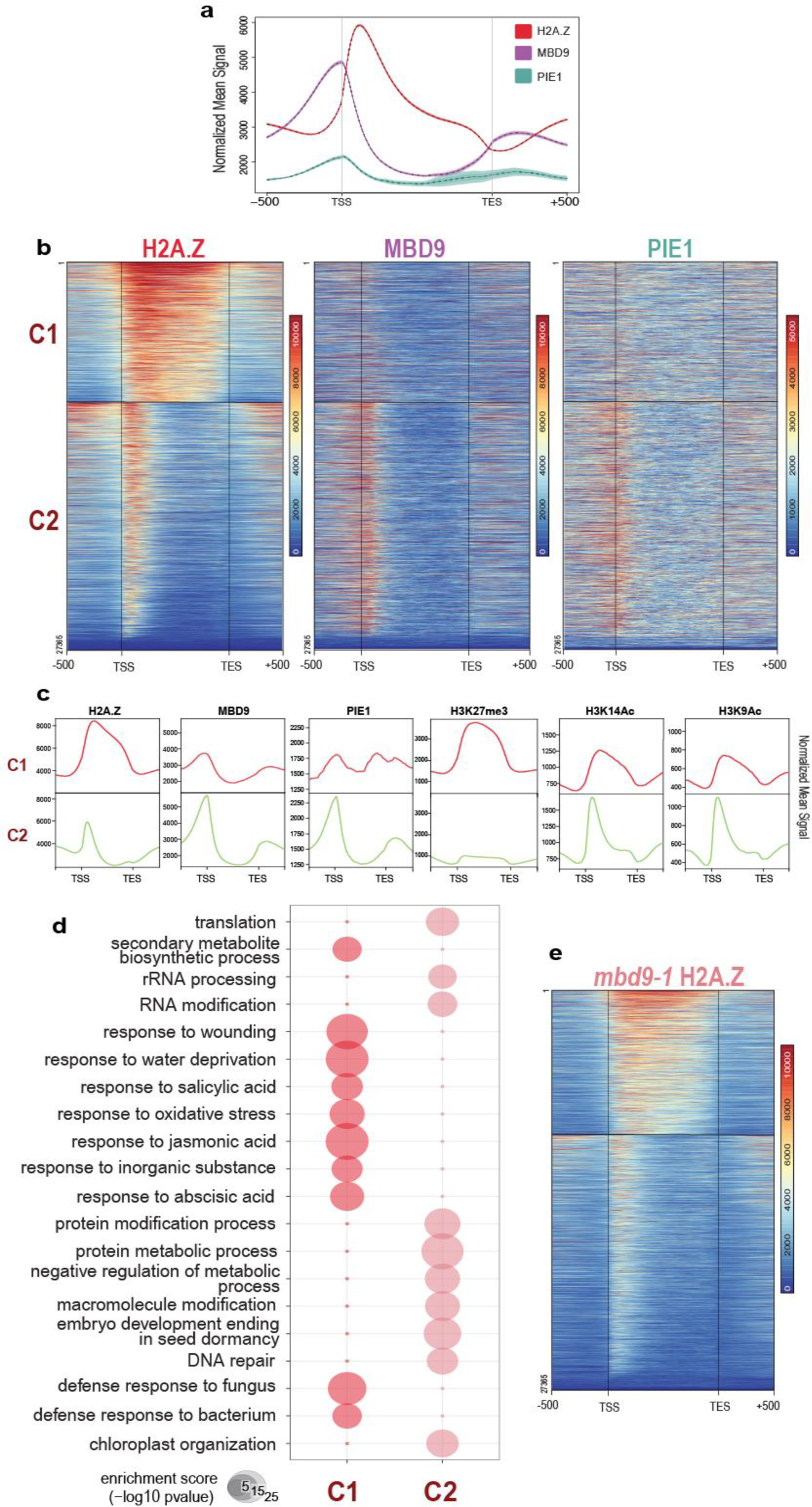
H2A.Z, MBD9 and PIE1 enrichment on genes at steady state. **(a)** Average profile plot showing enrichment of H2A.Z (red), MBD9 (purple) and PIE1 (teal) in WT plants on all Arabidopsis genes plotted from their Transcription Start Site (TSS) to their Transcription End Site (TES) and 500 base pairs up and downstream. The y axis indicates the enrichment of these proteins normalized to Reads Per Kilobase Million (RPKM). MBD9 and PIE1 enrichment are centered around the TSS while H2A.Z enrichment is highest at the nucleosome just downstream of the TSS (+1 nucleosome). **(b)** Representative heatmaps showing the enrichment of H2A.Z, MBD9 and PIE1 on all Arabidopsis genes broken into two k-means clusters (C1 and C2) based on WT H2A.Z. Locations are analogous across all heatmaps. Color saturation indicates the strength of H2A.Z, MBD9 or PIE1 at given genes plotted from TSS to TES. C1 has a higher enrichment of H2A.Z but less MBD9 and PIE1 compared to C2. Regions with high background omitted. Data are RPKM normalized. **(c)** Average profile plots showing the enrichment of H2A.Z, MBD9, PIE1, H3K27me3, H3K14Ac, and H3K9Ac on genes in WT plants at genes in C1(top row) or C2 (bottom row). C1 has more H2A.Z and H3K27me3 while C2 has more MBD9, PIE1, H3K14Ac and H3K9Ac comparatively. Patterns indicate genes in C1 are facultatively repressed while C2 encompasses actively transcribed genes. Data are RPKM normalized. PTM plots derived from publicly available data from analogous tissue (see Methods). **(d)** Bubble plot showing the top ten most enriched Gene Ontology (GO) categories (Fisher’s Exact Test, p-value <0.01) of C1 (red) and C2 (pink). Here, the size of each circle indicates the Enrichment Score (-log10 of p-value); the larger the circle, the more enriched the term. Only the top ten most enriched categories were shown here but there was no overlap in any significant terms across the two clusters. **(e)** Heatmap showing the enrichment of H2A.Z across C1 and C2 in *mbd9-1* plants. H2A.Z appears reduced in both C1 and C2 compared to WT.

When visualized with respect to all Arabidopsis coding genes, MBD9 and PIE1 localize mainly upstream of the transcription start site (TSS) in the nucleosome-depleted region (NDR), while H2A.Z enrichment begins at the first nucleosome downstream of the TSS (+1 nucleosome) (**Fig. 1a**). Genic H2A.Z enrichment begins at the +1 nucleosome and can occupy several nucleosomes downstream, but this enrichment skews to cover whole gene bodies in facultatively repressed genes ^16,23,29^. Genes therefore fall into one of two k-means clusters depending on their H2A.Z profile: gene body H2A.Z (gbH2A.Z) (**Fig. 1b**, ‘C1’) and TSS-proximal H2A.Z (**Fig. 1b**, ‘C2’). Interestingly, we see that C1 genes, which have the highest total H2A.Z levels relative to C2, have the least SWR1 (PIE1) and associated protein MBD9 (**Fig. 1b**). When investigating other histone PTMs associated with these two clusters, we see that C2 associates with PTMs implicated in active transcription, such as H3K14ac and H3K9ac while C1 is associated with H3K27me3, a PTM deposited by PRC2 that marks facultative heterochromatin (**Fig. 1c**). Gene ontology (GO) analysis shows that C1 has more significantly enriched terms associated with environmental responsiveness compared to C2 (**Fig. 1d**).

The lack of a strong correlation between H2A.Z abundance and PIE1 (SWR1) or MBD9 levels was surprising considering that PIE1 is directly involved in H2A.Z deposition and absence of MBD9 affects H2A.Z deposition in *mbd9-1* mutants (**Fig. 1e**). It is unclear if the distinction between C1 genes and C2 genes in reference to their patterning of H2A.Z, PIE1, and MBD9 are based on transcription status or are an inherent aspect of these genes.

### Transcriptional induction by abscisic Acid (ABA) causes subtle loss of H2A.Z and gains of PIE1 and MBD9

To understand how the enrichment of PIE1, MBD9, and H2A.Z change during widespread transcriptional alterations, we identified a subset of genes in C1 that could be inducibly activated. Although many groups of response genes are enriched in C1, genes implicated in the response to the plant hormone Abscisic Acid (ABA) were one of the significantly enriched GO pathways (**Fig. 1d**). These genes therefore represented a group whose transcription status could be altered with the exogenous application of ABA, providing us with a tool to study subsequent changes in H2A.Z, MBD9 and PIE1 localization. To this end, we treated 7-day-old WT whole seedlings with either 10 µM ABA or a Mock treatment for 4 hours and performed RNA-Seq and ChIP-Seq against H2A.Z, MBD9 and PIE1. To verify induction, we first performed RT-qPCR against two established ABA response genes, COR15A and ANAC72, and found that our treatment was sufficient to induce significant transcript increases in these genes in WT plants (**Supp. Fig. 2**). This significant change in transcript abundance was recapitulated in the RNA-Seq, where we measured 3417 Differentially Expressed Genes (DEGs) (1.5 fold change over Mock, padj<0.05) (**Fig. 2a**). Genes directly involved in ABA response were overrepresented among these DEGs (**Fig. 2b**). Although the DEGs encompassed genes in both C1 and C2, the mean change in transcript abundance of C1 genes was significantly higher than changes in C2, indicating that C1 genes are more responsive to ABA (Welch Two Sample t-test, p=1.72E-19)(**Fig. 2c**).

**Figure 2.**
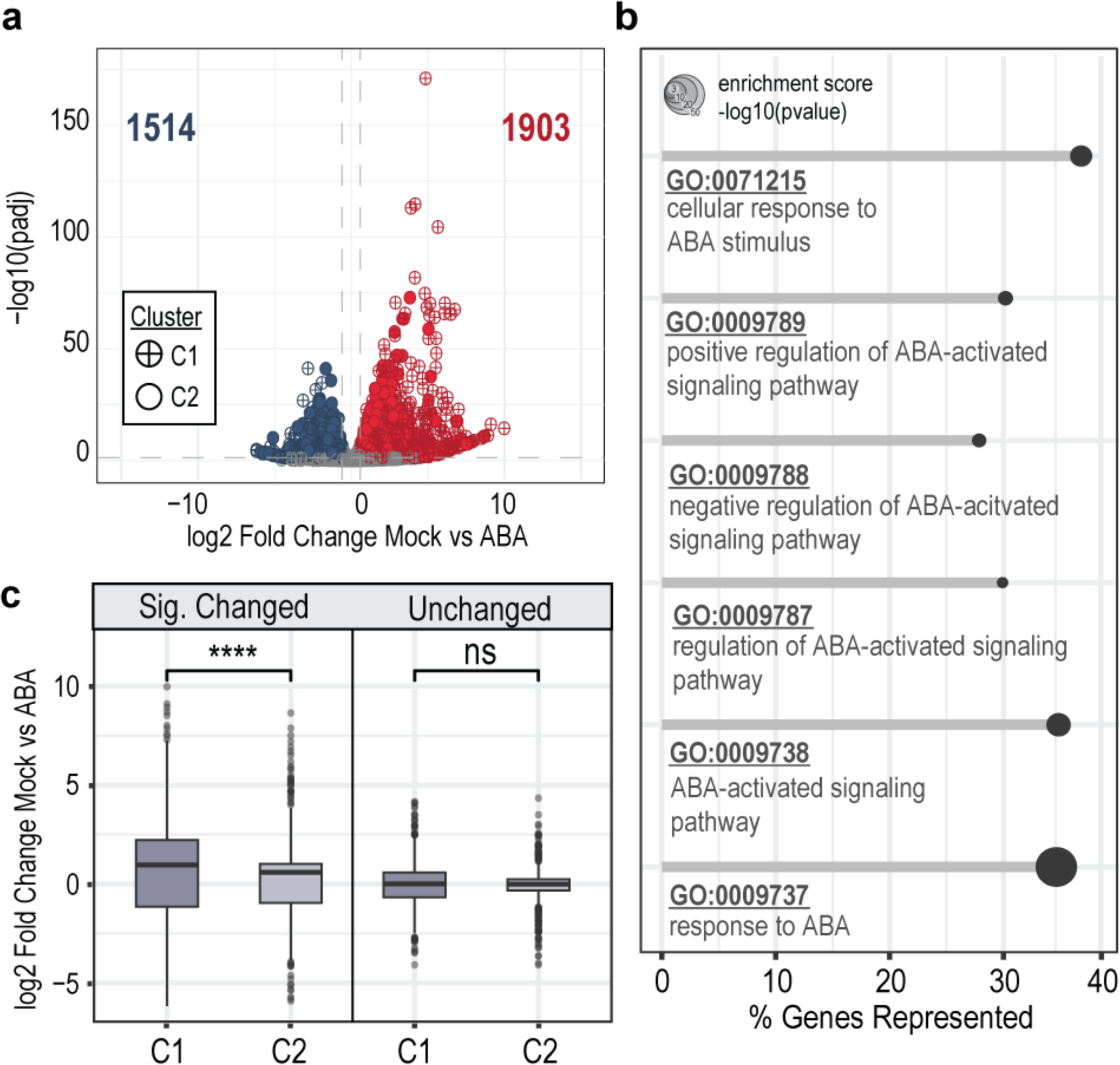
RNA-Seq changes in WT mock-treated plants vs WT ABA-treated plants. **(a)** Volcano plot showing all 3417 Differentially Expressed Genes (DEGs) in WT Mock vs ABA treated plants as measured by DESeq2. 1903 genes were Upregulated with a log2 Fold Change (log2FC) greater than 0.6 and an adjusted p-value less than 0.05 (red), while 1514 genes were Downregulated with a log2FC less than -0.6 and an adjusted p-value less than 0.05 (blue). Genes in Cluster 1 (C1) visually make up a greater proportion of the most significantly upregulated genes (crossed points). Dashed lines indicate p-value and log2FC significance thresholds. **(b)** Lollipop plot from GO Analysis of all DEGs shown in (a) displaying that several terms associated with ABA response are significantly enriched. GO names and brief descriptions are shown per category, and the length of the segment indicates the percentage of genes within that GO category represented in the 3417 DEGs. The size of the point indicates the enrichment score, calculated by taking the -log10 of the GO term p-value measured by Fisher’s Exact Test. **(c)** Boxplots depicting the log2FC of significantly changed DEGs or a random selection of Unchanged genes stratified by Cluster. The means of the Unchanged genes are not statistically different (Welch Two Sample t-test, t= 0.432, df = 1494, p=0.666) while the significantly changed DEGs are significantly different between clusters (Welch Two Sample t-test, t= 9.123, df = 2006, p=1.72E-19), reinforcing that C1 genes have more pronounced transcriptional changes in response to ABA compared to C2 in WT plants.

ChIP-sequencing data also revealed minor changes in H2A.Z abundance and larger changes in PIE1 and MBD9 abundance at the 1342 C1 DEGs after ABA treatment. These changes are most evident at the 813 C1 Upregulated genes (**Fig. 3a-3c**, middle panel), where H2A.Z levels show a uniform but minor decrease after ABA treatment, with a simultaneous gain in MBD9 and PIE1 enrichment. The inverse is true at the 529 C1 Downregulated genes, where there is a slight gain in H2A.Z and a subtle loss of MBD9 and PIE1 in ABA-treated plants (**Fig. 3a-3c**, top panel). Importantly, these subtle changes were not observed in an equivalently-sized set of C1 genes whose expression was not changed after ABA treatment (**Fig. 3a-3c**, bottom panel). These changes can also be observed on average plots of these three proteins across the different gene groups (**Fig. 3d**). Here we focus on C1 genes, but similar patterns are observed at C2 genes. See **Supp Fig. 3** for equivalent heatmaps showing the dynamics at C2 DEGs.

**Figure 3.**
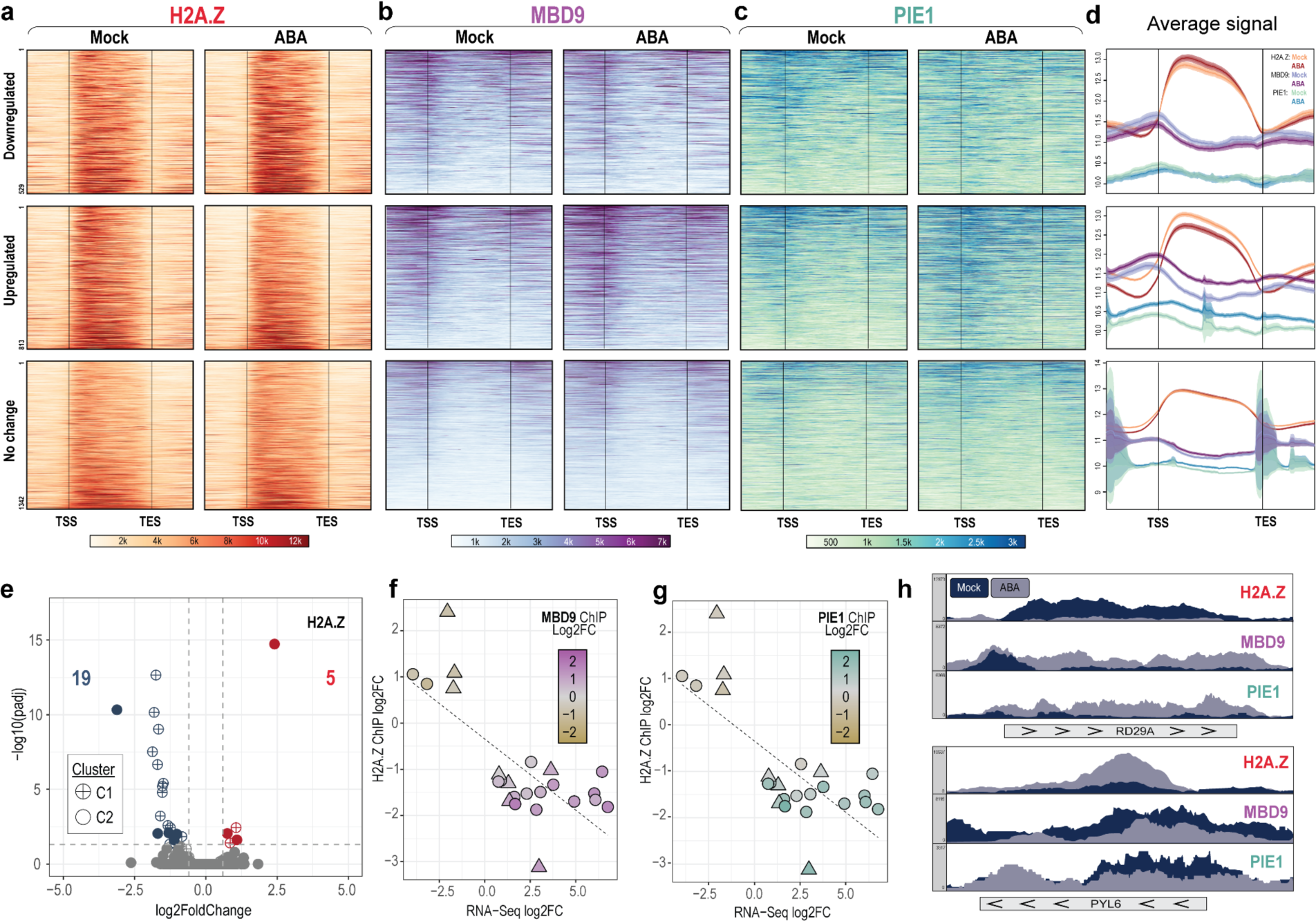
C1 DEGs show losses of H2A.Z and gains in MBD9 and PIE1 proportional to induction. **(a)** Representative heatmaps showing H2A.Z enrichment on 529 Downregulated DEGs (Top panel), 813 Upregulated DEGs (Middle panel) and 1342 C1 genes with no significant change in transcript abundance after ABA treatment (Bottom panel). Enrichment is plotted from the Transcription Start Site (TSS) to the Transcription End Site (TES). At Upregulated genes, there is a loss of H2A.Z enrichment and a gain of MBD9 and PIE1 as shown in similar representative heatmaps in **(b)** and (**c).** All data are RPKM normalized. **(d)** Average plots showing the average signal of all genes at Downregulated (Top panel), Upregulated (Middle panel), and genes with No Change (Bottom panel). RPKM Signal from all genes is averaged and plotted on a log2 scale from their TSS to TES. **(e)** Volcano plot showing 24 genes differentially enriched in H2A.Z as measured by DESeq2 in Mock vs ABA treated plants. 16 genes belong to C1 (crossed points) while 8 belong to C2 (solid points). Blue points show the 19 genes that lost H2A.Z with ABA treatment (log2FC < -0.6 and adjusted p-value<0.05) and red points show the 5 genes that gained H2A.Z with ABA treatment (log2FC>0.6 and an adjusted p-value< 0.05, red points). 14 and 2 of the H2A.Z lost and H2A.Z gained genes belonged to the C1 cluster, respectively. **(f)** 24 genes with differential H2A.Z enrichment (y-axis) correlated with RNA-Seq expression (x-axis) and enrichment of MBD9 (color). There is a negative correlation between H2A.Z and transcript changes in Mock vs ABA treated plants. Genes that lose H2A.Z and increase in transcript also have the highest gain of MBD9. **(g)** shows the same trends with PIE1 changes.**(h)** Integrative Genomics Viewer (IGV) snapshots of eChIP-Seq data at two genes illustrating patterns of H2A.Z, MBD9 and PIE1 changes. *RD29A (AT5G52310)* shows a loss of H2A.Z and a gain of MBD9 and PIE1 in ABA treated plants while the *PYL6 (AT2G40330)* gene shows a gain of H2A.Z and a loss of MBD9 and PIE1 signal. Both genes belong to Cluster 1 (C1).

Surprisingly, there were only 24 genes genome wide that had a statistically significant change in H2A.Z after ABA treatment as measured by DESeq2, and 16 of these were in C1 (adjusted p-value<0.05 and log2FC < -0.6 or >0.6) (**Fig. 3e**). More of the genes that lost H2A.Z belonged to C1 (14/19), and almost all of the genes that gained H2A.Z belonged to C2 (3/5). When layering our RNA-Seq and ChIP-Seq data, we found a correlation between H2A.Z change, transcript abundance, and PIE1 and MBD9 change upon ABA treatment (**Fig. 3f and g**). Of genes with differential H2A.Z enrichment after ABA treatment, the genes with gains in transcript correlated with loss of H2A.Z, and the genes with loss of transcript correlated with gain of H2A.Z. When MBD9 and PIE1 are considered, we also see that they are recruited most strongly to the genes that gain transcript and lose H2A.Z (**Fig. 3f and g**). This pattern is also observed on a gene-by-gene basis when considering two of the genes with significantly changed H2A.Z (**Fig. 3h**). These correlations suggest that the level of H2A.Z is a reflection of the balance between nucleosome disruption by transcription machinery and subsequent re-incorporation of H2A.Z by SWR1 at these inducible genes, with MBD9 potentially playing a role in this process. It is also notable that the original gene body enrichment of H2A.Z on upregulated C1 genes is maintained after transcriptional change. The abundance of H2A.Z increases or decreases, but it is still localized across the gene body and does not convert to TSS-proximal H2A.Z enrichment (**Fig. 3a-d, middle panels**).

### The MBD9 Bromodomain is Dispensable for proper H2A.Z Localization

Based on these observations, we hypothesized that MBD9 is recruited to newly activated ABA responsive genes through its Bromodomain, which has been shown previously to associate with acetylated histone residues characteristic of active transcription including H3K14Ac and H3K18Ac ^47^. To assess the contribution of the MBD9 Bromodomain, we aimed to specifically disrupt the Bromodomain function without affecting other domains. MBD9 has several putative domains other than its acetyl-lysine binding Bromodomain including two methyl-binding Plant HomeoDomains (PHD), and a Williams-Beuren Syndrome DDT (WSD) domain that may mediate interactions with Imitation SWItch (ISWI) type chromatin remodelers (**Fig. 4a**) ^48,50^. MBD9 is also one of 13 family members in Arabidopsis that contain a Methyl-CpG Binding Domain (MBD). Normally, this domain facilitates interactions with methylated DNA, but MBD9 contains several point mutations in its MBD sequence that prevent this function ^51,52^.

**Figure 4.**
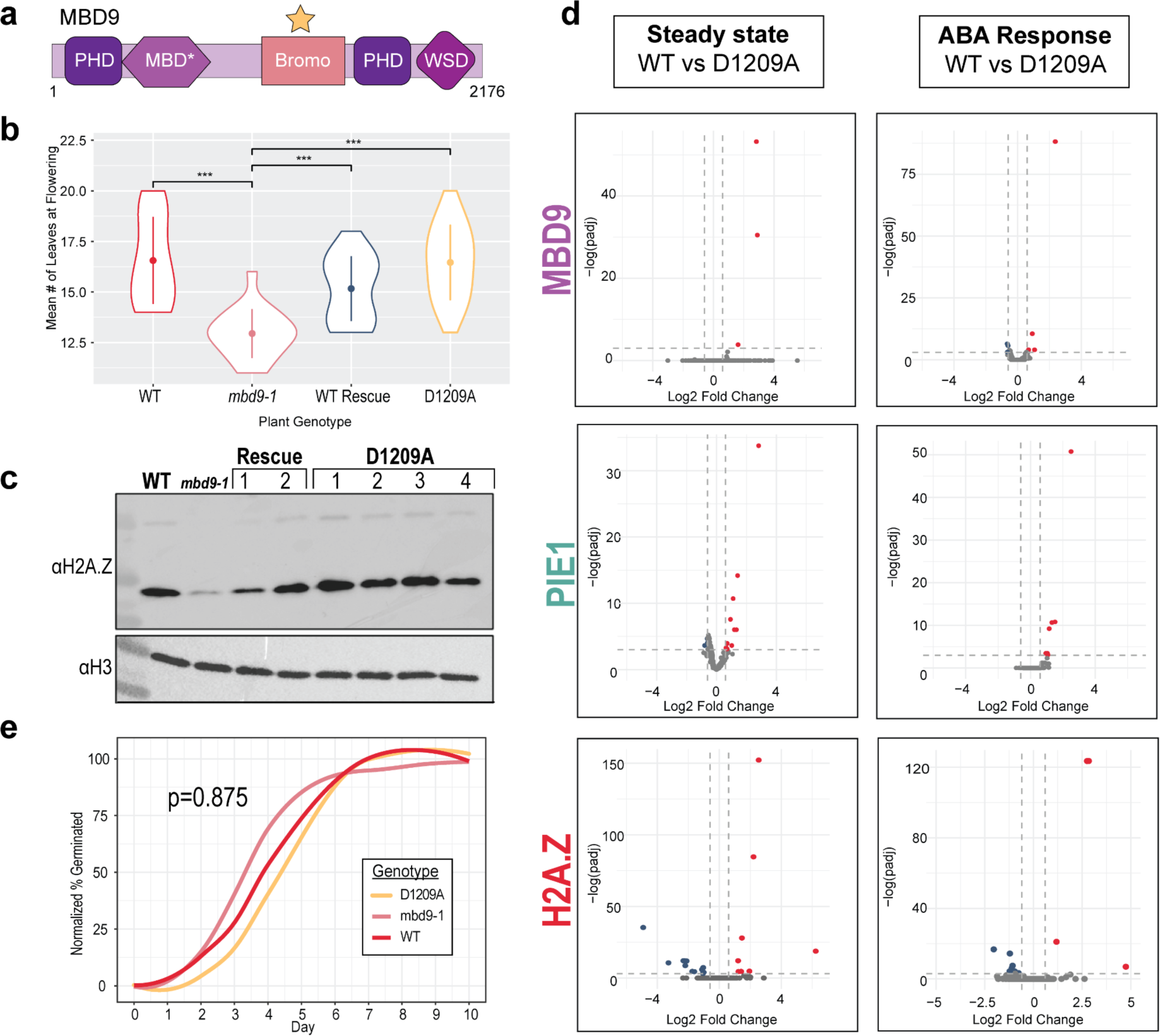
D1209A mutants at steady state and during an ABA Response. **(a)** Diagram of MBD9 showing positioning of domains including Plant HomeoDomains (PHD), Bromodomain (Bromo), Williams-Beuren Syndrome DDT (WSD) domain, and Methyl-Binding Domain (MBD). Organization not to scale. *MBD has point mutations preventing binding to methylated DNA. **(b)** Violin plot showing flowering time of D1209A plants in reference to WT, plants lacking MBD9 (*mbd9-1*), and plants rescued with WT MBD9 (WT Rescue). *mbd9-1* plants flower early compared to WT plants (Welch Two Sample t-test, t=-6.2125, df= 26.819,p=1.25E-06). Supplementation with WT MBD9 rescues early flowering (Welch Two Sample t-test, t=-5.1115, df= 39.994,p=8.30E-06), and D1209A plants rescue early flowering (Welch Two Sample t-test, t=9.0273, df= 46.365, p=8.84E-12). Means for WT Rescue and D1209A plants represent average across two or four independent insertion lines, respectively. ***= p-value<0.001**(c)** Western blot of chromatin associated proteins for four independent D1209A lines show more H2A.Z compared to *mbd9-1*. Two independent WT Rescue lines also show more H2A.Z incorporation compared to *mbd9-1*. H3 shown as a loading control. **(d)** Volcano plots showing differences in the localization of MBD9 (top), PIE1 (middle) and H2A.Z (bottom) in WT vs D1209A plants. Blue points are genes with less H2A.Z in D1209A plants (log2 Fold Change< -0.6, adjusted p-value<0.05) and red points are genes that gain H2A.Z in D1209A plants (log2 Fold Change>0.6, adjusted p-value< 0.05). Plots on the left show differences at steady state while plots on the right are during an ABA response. There are few differentially enriched genes in any of these conditions. **(e)** Line graph depicting the germination efficiency of D1209A plants on ABA-containing media compared to WT and *mbd9-1* plants over time. The number of germinated seeds is expressed as a percentage of total seeds over the course of 10 days, normalized to the genotype germination on unsupplemented ½ MS media. Average across two independent germination experiments shown. There was no significant effect of genotype on germination efficiency (ANOVA, f=0.134, df=2, p=0.875).

Aspartic acid 1209 of MBD9 is a well-conserved residue that spans various bromodomain-containing proteins (**Supp. Fig. 4**), and was previously shown to be necessary for its acetylated histone peptide binding function *in vitro* ^47^. We mutated this Aspartic Acid 1209 to an Alanine (D1209A) in the MBD9 sequence through site-directed mutagenesis and established stable Arabidopsis lines expressing D1209A MBD9 in the null *mbd9-1* background (D1209A plants). At the phenotypic level, we see that D1209A plants do not show an early flowering phenotype characteristic of plants lacking MBD9. D1209A plants had significantly more leaves at flowering compared to *mbd9-1* plants (Welch Two Sample t-test, p=8.84E-12) (**Fig. 4b**). Western blotting of chromatin-associated proteins from 7-day-old D1209A plants also demonstrates that D1209A MBD9 rescues the ability of H2A.Z to be incorporated into chromatin compared to *mbd9-1* plants (**Fig. 4c**). ChIP-Seq against MBD9, PIE1 and H2A.Z in D1209A plants at steady state and after treatment with ABA reveal that the vast majority of MBD9, PIE1 and H2A.Z sites were unchanged from WT as measured by DESeq2 both at steady-state and during an ABA response (**Fig. 4d**). D1209A plants also show no significant difference in germination rate on ABA-containing media compared to WT and *mbd9-1* plants (**Fig. 4e**). Taken together, these results demonstrate that the Bromodomain of MBD9 is dispensable for its effect on H2A.Z localization.

### Loss of MBD9 does not affect SWR1 recruitment or transcriptional ABA responsiveness

To further assess if MBD9 is required for ABA-driven changes in transcription, we also performed RNA-Seq and ChIP-seq against PIE1 and H2A.Z in Mock and ABA treated plants completely lacking MBD9 (*mbd9-1*; **Supp Fig 1**). Comparison of Mock treated WT and *mbd9-1* plants revealed that only 184 genes are differentially expressed on a genotype basis alone (**Fig. 5a**). *mbd9-1* plants have a robust transcriptional response to ABA, with 1950 genes being Upregulated and 1716 genes being Downregulated compared to Mock Treatment (**Fig. 5b**). Overall, 75% of *mbd9-1* significantly Upregulated genes and 59% of *mbd9-1* significantly Downregulated genes are conserved across genotypes in response to ABA (**Fig. 5c**). Although there are ‘uniquely’ Upregulated and Downregulated genes specific to each genotype, direct correlation of changes at these genes reveals that *mbd9-1* and WT responses trend similarly but do not reach statistical significance in one genotype (**Supp Fig 5**). Focusing on WT DEGs, we see that *mbd9-1* and WT changes in response to ABA are strongly correlated, but with WT changes being of slightly higher magnitude in both the negative and positive directions (**Fig. 5d**). This trend is evident at both C1 and C2 genes separately, though the mean change between clusters is not statistically significant (**Fig. 5e**).

**Figure 5.**
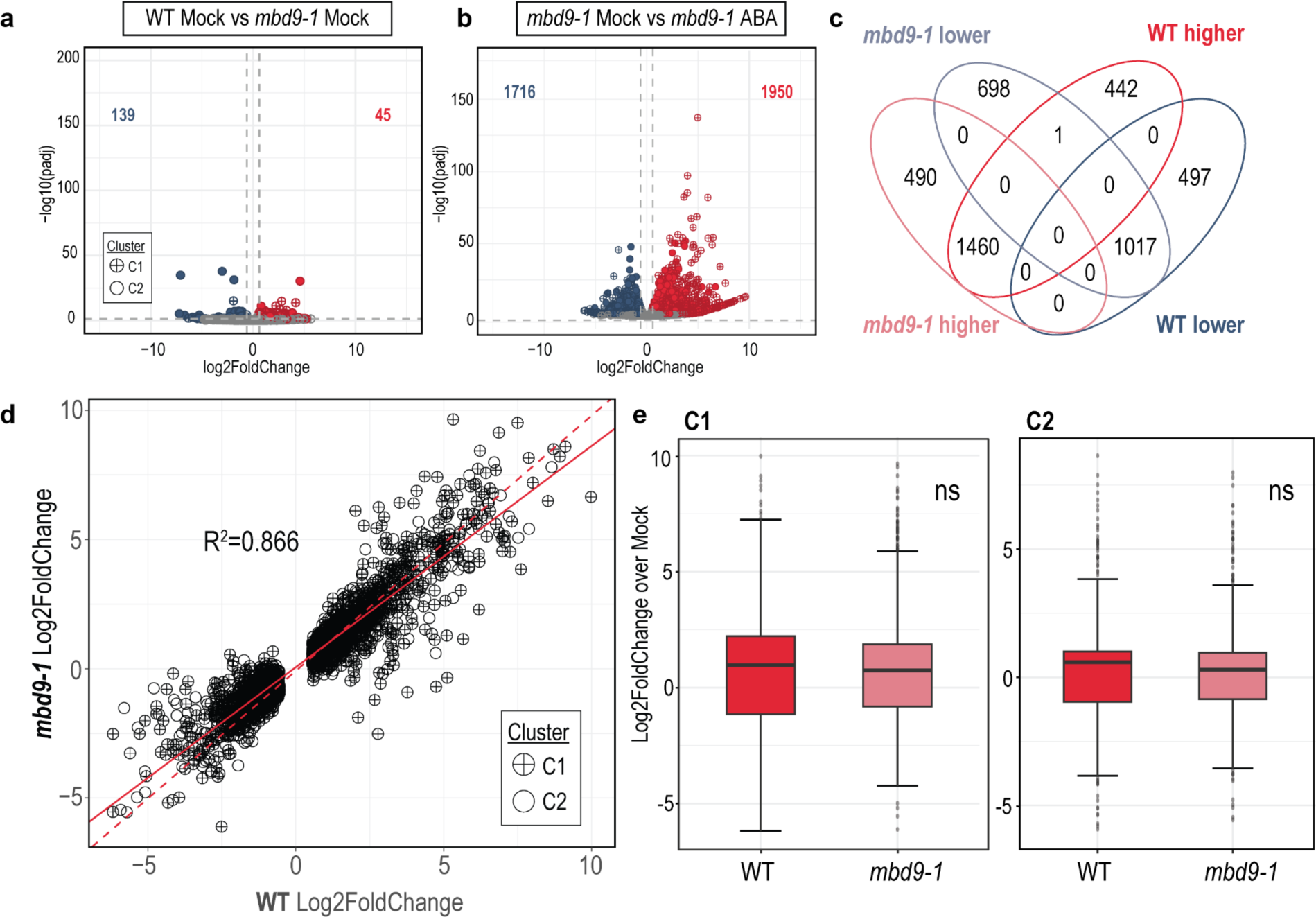
*mbd9-1* RNA responses compared to WT RNA responses. **(a)** Volcano plot showing differentially expressed genes (DEGs) in WT vs *mbd9-1* Mock treated plants, identifying genes that are differentially expressed on a genotype basis. Blue points show genes downregulated in *mbd9-1* (log2 Fold Change< -0.6, adjusted p-value<0.05) while red points show genes upregulated in *mbd9-1* (log2 Fold Change>0.6, adjusted p-value< 0.05, red points). There are 45 and 139 Downregulated and Upregulated genes, respectively. **(b)** Volcano plot showing differentially expressed genes (DEGs) in mbd9-1 Mock vs *mbd9-1* ABA treated plants. *mbd9-1* shows 1716 downregulated genes (log2 Fold Change< -0.6, adjusted p-value<0.05) and 1950 upregulated genes (log2 Fold Change>0.6, adjusted p-value< 0.05). **(c)** Direct comparison of DEGs in WT plants vs *mbd9-1* plants. The majority of up and downregulated genes are conserved across genotypes, but unique subsets exist in each genotype. See Supp Fig. 5 for more info on these groups.**(d)** Scatter plot correlating RNA-Seq Log2 FoldChanges between WT and *mbd9-1* plants at WT DEGs during ABA response. The dashed red line indicates a hypothetical 1:1 relationship between WT and *mbd9-1* responses while the solid red line indicates the linear regression of the data. The R^2^ value between the groups is 0.866, indicating a strong relationship between WT and *mbd9-1* response. C1 genes are denoted by crossed points whereas C2 genes are denoted by uncrossed points. **(e)** Boxplots depicting log2FoldChanges of RNA changes in ABA vs Mock treated samples across WT and *mbd9-1* plants, stratified by Cluster. Cluster 1 (left) genes show slightly tempered changes compared to WT plants, but these changes do not reach statistical significance (Welch Two Sample t-test, t=0.21, df=2665, p=0.835 ). The same is observed at Cluster 2 (right) (Welch Two Sample t-test, t=-0.35, df=4111, p=0.725).

At the chromatin level, we see that *mbd9-1* plants, although they start with less H2A.Z at genes than WT (compare **Fig. 1b and 1e**), still lose H2A.Z from ABA Upregulated genes as in WT plants (**Fig. 6a**, middle panel). Similarly, the ABA Downregulated genes gain H2A.Z in ABA-treated *mbd9-1* plants (**Fig. 6a**, top panel). Also similar to WT, PIE1 levels increase specifically at ABA Upregulated genes in *mbd9-1* (**Fig. 6b**, middle panel). These changes are also evident on average (**Fig. 6c**). When directly comparing average PIE1 changes at ABA upregulated genes, it is apparent that while *mbd9-1* plants may start with slightly less PIE1 at upregulated genes, the gain in PIE1 during ABA treatment is equivalent to gains in WT plants (**Fig 6d**). Compared to only 24 genes in WT, *mbd9-1* shows 220 genes differentially enriched in H2A.Z after ABA treatment compared to Mock treatment, with the genes losing the most H2A.Z belonging to C1 (**Fig. 6e**). When correlating the significant chromatin changes with RNA-Seq data, we see the genes that gain the most transcript lose the most H2A.Z and gain the most PIE1 in *mbd9-1* plants (**Fig. 6f**), as observed in WT plants.

**Figure 6.**
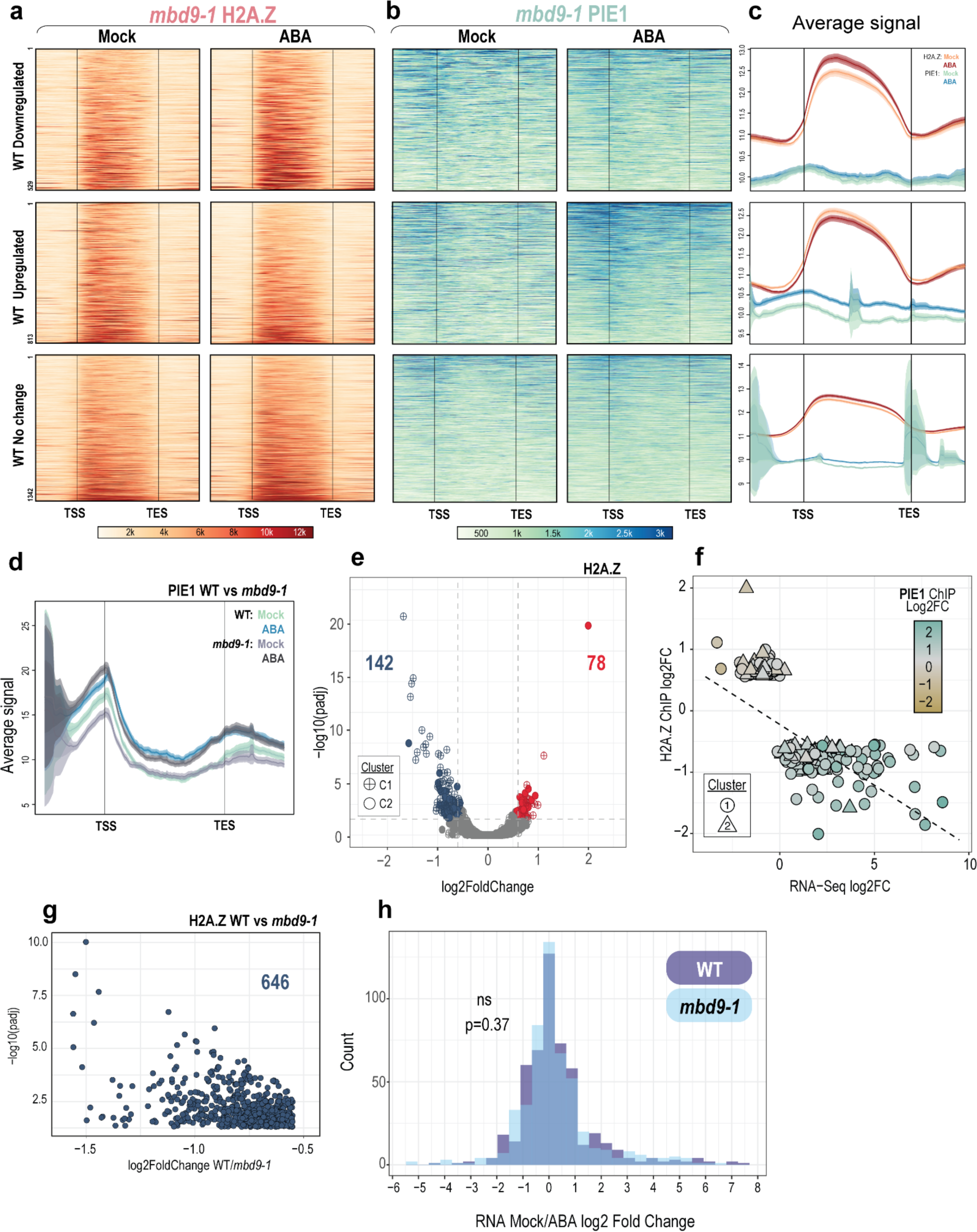
ABA-treated *mbd9-1* plants show similar trends in chromatin dynamics. **(a)** Representative heatmaps showing RPKM normalized H2A.Z enrichment on genes from Fig. 3: 529 Downregulated (Top panel), 813 Upregulated (Middle panel) and 1342 C1 genes with no significant transcript change after ABA treatment (Bottom panel). Enrichment plotted from the Transcription Start Site (TSS) to the Transcription End Site (TES). *mbd9-1* plants show a similar trend, where Upregulated genes lose H2A.Z after ABA treatment and downregulated genes gain H2A.Z.**(b)** Representative heatmaps of PIE1 enrichment over the same areas as **(a)**. PIE1 signal still increases after ABA treatment at Upregulated genes (middle panel).**(c)** Average plots depicting H2A.Z and PIE1 signal at genes that are Downregulated (Top panel), Upregulated (Middle panel), and genes with No Change (Bottom panel). RPKM Signal from all genes is averaged and plotted on a log2 scale from TSS to TES. **(d)** Average plots directly comparing average PIE1 changes at Upregulated genes in WT vs *mbd9-1* plants. *mbd9-1* PIE1 signal at upregulated genes is roughly equivalent in ABA treated conditions. RPKM Signal from all genes is averaged and plotted on a log2 scale from TSS to TES. **(e)** Volcano plot showing 220 genes differentially enriched in H2A.Z as measured by DESeq2 in Mock vs ABA treated *mbd9-1* plants. 142 genes lose H2A.Z (log2 Fold Change< -0.6, adjusted p-value<0.05, blue) and 78 genes gain H2A.Z with ABA treatment (log2 Fold Change>0.6, adjusted p-value< 0.05, red). **(f)** 220 genes with differential H2A.Z (y-axis) correlated with RNA-Seq expression (x-axis) and abundance of PIE1 (color). The negative correlation between H2A.Z and transcription and positive correlation with PIE1 is maintained in *mbd9-1*. **(g)** Scatter plot showing 646 genes with lower H2A.Z in *mbd9-1* plants compared to WT (padj<0.05, log2FoldChange<-0.6). **(h)** Histogram showing that the 646 genes from **(g)** change similarly in transcript abundance over Mock in WT (purple) vs *mbd9-1* plants in response to ABA (Student’s t-test, t=0.89904, df= 1010, p=0.3688), indicating that although these genes have less H2A.Z to start with in *mbd9-1* plants, it does not affect their responsiveness during a stress response.

These data suggest that transcription rate determines the amount of H2A.Z lost, and not vice versa. To more directly rule out the possibility that the starting amount of H2A.Z affects the induction potential of genes, we isolated a set of 646 genes that have significantly less H2A.Z in *mbd9-1* Mock treated plants compared to WT Mock treated plants (**Fig. 6g**). If the amount of H2A.Z affects the induction or repression of these genes, we should see that transcript levels in *mbd9-1* plants, which have significantly less H2A.Z at these genes, should change less substantially after ABA treatment compared to WT. Instead, we see that the distributions of log2 Fold Changes of these genes’ transcripts are not statistically different after ABA treatment, supporting the notion that the reduced amount of H2A.Z does not affect the induction potential of these genes (**Fig. 6h**).

## Discussion

By utilizing new antibodies against SWR1 component PIE1 and associated protein MBD9, we examined the relationships between H2A.Z, MBD9, SWR1, and transcription in plants. We demonstrated that at steady state, the genes most enriched with H2A.Z (C1) have the least amount of SWR1 (PIE1) and MBD9 binding (**Fig 1a-1c**). One explanation for this phenomenon is simply that C1 genes, which encompass genes marked with H3K27me3, are not heavily transcribed under standard physiological conditions. Thus, without transcription machinery consistently disrupting these genic, stable nucleosomes, nucleosome turnover is low and there is no impetus for SWR1 to replace the H2A.Z-containing nucleosomes lost during transcription. At the more constitutively expressed C2 genes, which have TSS-proximal H2A.Z, nucleosomes are more unstable and require sustained presence of SWR1 to maintain H2A.Z enrichment. Transcript count data from Mock treated plants here and publicly available Global Run-on Sequencing (Gro-Seq) data from analogous tissue support this hypothesis ^53^; C1 genes have significantly fewer counts than C2 genes in these datasets (**Supp. Fig. 6**).

To understand how PIE1 and MBD9 localizations change after activation of genes with gbH2A.Z (C1 genes), we employed ABA as a widespread transcriptional modulator to assess if changes to transcription in a subset of C1 genes led to altered recruitment of core SWR1 component PIE1 and known interacting protein MBD9. In WT plants, we saw that the genes with the highest gains in transcript abundance had the largest loss of H2A.Z and highest gains of MBD9 and PIE1 (**Fig. 3f and 3g**). This suggests that when these formerly silenced genes are activated transcriptionally, the SWR1 complex is recruited to replace the H2A.Z lost by transcription, thereby maintaining H2A.Z presence on these genes.

While the majority of H2A.Z at ABA response genes appeared unchanged after induction, it is likely that these ‘unchanged’ genes underwent constant transcription-driven loss and SWR1-driven replacement of H2A.Z. We hypothesize that at these unchanged sites, SWR1 deposited H2A.Z at a rate closely matching the rate of H2A.Z loss caused by transcription. However, at the genes with the largest upregulation of transcription, the SWR1 complex could not keep up with the rate of H2A.Z dissociation from genes and these 19 genes significantly lost H2A.Z (**Fig. 3e**). This hypothesized high turnover of H2A.Z could then also result in exchange of repressive monoubiquitinated H2A.Z for unmodified H2A.Z by SWR1, potentially contributing to depression of these genes ^29^. The phenomenon of gbH2A.Z retention during transcriptional activation of silent genes was also recently observed for phosphate-starvation induced genes in Arabidopsis root hair cells (Holder and Deal 2024 ^81^). These results indicate that net H2A.Z removal from chromatin is not strictly required for transcriptional activation. This is in contrast to previous studies showing that H2A.Z removal by the chromatin remodeler INO80 is necessary for induction of environmental response genes including those responsive to temperature, far-red light, and ethylene ^38–40,54^. Collectively, these results suggest the existence of both INO80-dependent and -independent pathways for activation of silent gbH2A.Z/H3K27me3 genes. While some genes may require H2A.Z removal for activation, others such as ABA responsive genes instead seem to require SWR1 recruitment to retain gbH2A.Z.

Our data also showed that MBD9, which was previously shown to interact with the SWR1 complex, was also recruited to newly activated genes. Based on this observation, we asked whether the Bromodomain of MBD9 was important for this recruitment, and perhaps that of SWR1. We used a mutation that was shown previously to hinder the ability of the MBD9 Bromodomain to bind acetylated histone peptides *in vitro* ^47^, but demonstrated that this mutation has little effect on plants *in vivo*. We saw that the D1209A Bromodomain mutant MBD9 can rescue the early flowering phenotype of *mbd9-1* plants (**Fig. 4b**). This recapitulated flowering time experiments performed on Bromodomain mutants previously, where Arabidopsis *mbd9-1* plants supplemented with an MBD9 harboring the D1209A mutation in combination with E1208A showed rescue of *mbd9-1* early flowering ^48^. We also found that at the molecular level, this mutation had little effect on the localization of MBD9, PIE1 or H2A.Z on genes either at steady state or when responding to ABA (**Fig. 4d**). If the Bromodomain is truly dispensable for the role of MBD9 in H2A.Z localization, it is in contradiction to a previous model that posits MBD9, binds to acetyl marks through its Bromodomain as a way to anchor the SWR1 complex for H2A.Z deposition ^47^. This model, however, also suggests that another bromodomain-containing protein NPX1, which is partially conserved with yeast BDF1, acts redundantly with MBD9 in this SWR1 anchoring. NPX1 may be able to compensate for MBD9 Bromodomain loss in the context of this study. Alternatively, the MBD9 Bromodomain could mediate interactions with acetylated histones but this is not relevant to H2A.Z localization. Future work with MBD9 will uncover whether other domains are necessary for its role in proper H2A.Z localization and explore the role of the MBD9 Bromodomain in functions outside of the SWR1 complex.

To further assess the role of MBD9 during transcriptional activation, we determined how plants lacking MBD9 (*mbd9-1*) responded to ABA. Under steady state, it was previously reported that *mbd9-1* plants do not have widespread de-repression of response genes ^50^. Our results support this as well; we measured only 184 genes as differentially expressed in *mbd9-1* plants compared to WT on a genotype basis alone (**Fig. 5a**). We also demonstrated that *mbd9-1* plants effectively responded to ABA, with the majority of genes induced similarly in *mbd9-1* compared to WT (**Fig. 5d**). While loss of MBD9 did not drastically affect plants’ transcriptional response to ABA, trends at both C1 and C2 genes suggest that *mbd9-1* plants may not be able to induce or repress genes as effectively as in WT plants (**Fig. 5e**). These subtle changes in transcriptome response could be attributable to the reported role of MBD9 in the ISWI complex ^48^. MBD9 likely serves as a bridge between remodelers, so without its presence, ISWI would not be recruited to newly activated genes to help retain newly deposited H2A.Z in chromatin. This could lead to the subtle changes in transcription dynamics of these genes with high nucleosome turnover after induction.

At the chromatin level, *mbd9-1* plants behaved similarly to WT in regard to H2A.Z and PIE1 localization. Although *mbd9-1* plants had less H2A.Z at genes on average, there was still a subtle loss of H2A.Z from upregulated genes, and a subtle gain of H2A.Z at downregulated genes (**Fig. 6a**). Interestingly, the absence of MBD9 did not inhibit SWR1 recruitment to ABA-upregulated genes (**Fig. 6b**), indicating that it is not the major targeting factor for SWR1 in this context. This is further reinforced by direct comparison of PIE1 changes in WT and *mbd9-1* plants; PIE1 was gained in ABA treated *mbd9-1* plants to the same degree as in WT (**Fig. 6d**). Instead, specific transcription factors or even the transcription machinery could directly recruit chromatin remodelers such as SWR1 to newly activated genes. A recent study demonstrated that the BAF complex is recruited by RNA Pol II upon transcriptional change in mouse embryonic stem cells ^55^. It is also interesting to note that *mbd9-1* plants, which have less H2A.Z on average, did not show widespread differences in gene expression upon ABA activation compared to WT plants (**Fig. 6h**). This indicates that while H2A.Z contributes to a repressive state, the degree of H2A.Z loss in *mbd9-1* is not sufficient to derepress genes or change their responsiveness to a great extent.

We also saw that the profiles of gbH2A.Z were actively maintained in the face of transcriptional activation by ABA. It has been proposed that the two distinct profiles of H2A.Z, TSS-proximal H2A.Z (C2) and gbH2A.Z (C1) may be interconvertible. Genes with gene body H2A.Z, once activated, will shift to having TSS-proximal H2A.Z enrichment, and then revert to having gbH2A.Z after repression is re-established ^27,56,57^. Here, however, changes to H2A.Z within each cluster reflected the original profile. C1 genes with lower H2A.Z after ABA treatment lost H2A.Z across the gene body, and genes with higher H2A.Z gained H2A.Z over the gene body (**Fig. 3d and Fig. 6c**). Similarly, C2 genes both gained and lost H2A.Z at TSS-proximal nucleosomes after ABA treatment on average (**Supp. Fig 3**). These results are consistent with similar studies analyzing H2A.Z distribution in Arabidopsis during drought stress ^27^. Clearly the cell is utilizing energy to maintain these two distinct profiles of H2A.Z, possibly highlighting that TSS-proximal H2A.Z and gbH2A.Z are distinct pools with separate functions that do not reflect current transcription rate as accurately as previously thought.

In summary, by tracking SWR1 component PIE1 and interactor MBD9, we gained new insights into the roles of these proteins in H2A.Z maintenance during a widespread transcriptional activation event. Surprisingly, at steady state we saw that genes with the most H2A.Z had the least amount of SWR1 component PIE1 and interactor MBD9. After treating plants with ABA, we saw that PIE1 and MBD9 move to newly induced genes, with the genes losing the most H2A.Z gaining the most PIE1 and MBD9. This indicates that SWR1 moved to activated genes to replace H2A.Z-containing nucleosomes lost by transcription. Although MBD9 was recruited to newly activated genes, its presence was largely not necessary for PIE1 recruitment or a normal transcriptional response to ABA. This indicates that MBD9 does not serve as a general SWR1 targeting factor, but instead may be necessary for full functionality of the SWR1 complex or could affect H2A.Z occupancy indirectly through ISWI-related functions.

## Methods

### Plant growth conditions

All *Arabidopsis thaliana* plants described are of the Columbia (Col-0) ecotype, and Col-0 was used as a wild type (WT) reference. Described mutants obtained from ABRC include *mbd9-1* (SALK_054659) and *pie1-5* (SALK_053834C). Seeds were sown on half-strength Murashige and Skoog (MS) media plates or soil and stratified at 4°C for 72 hours. After stratification, plants were moved to growth chambers at 20°C under a 16 hour light/8 hour dark cycle under 181 µmol m^-2^ s^-1^ PPFD light intensity. Unless otherwise noted, plated plants were grown approximately 7 days and whole seedling shoot tissue (above ground tissue only) was collected at approximately 9 hours after artificial dawn for use in downstream experiments.

### Antibody Development

Polyclonal antibodies against Arabidopsis proteins PIE1 (AT3G12810) and MBD9 (AT3G01460) were made using services from GenScript. The MBD9 protein fragment ‘MEPSILKEVGEPHNSSYFADQMGCDPQPQEGVGDGVTRDDETSSTAYLNKNQGKSPLETDTQPGESHVNFGESKISSPETISSPGRHELPIADTSPLVTDNLPEKDTSETLLKSVGRNHETHSPNSNAVELPTAHDASSQASQELQACQQDLSATSNEIQNLQQSIRSIESQLLKQSIRRDFLGTDASGRLYWGCCFPDENPRILVDGSISLQKPVQADLIGSKVPSPFLHTVDHGRLRLSPWTYYETETEISELVQWLHDDDLKERDLRESILWWKRLRYGDVQKEKKQAQNLSAHHHHHH’ was expressed recombinantly and the PIE1 peptide fragment ‘CEEIRKAVFEERIQESKDRAAAI’ was synthesized for use as antigens in polyclonal antibody production in rabbits. The antibodies were antigen affinity purified by the manufacturer and verified here through Western Blotting (**Supp. Fig. 1**). Additionally, an identical MBD9 peptide was used as an antigen for antibody production previously ^58^.

### MBD9 BromoDomain (D1209A) Mutant Generation

The genomic sequence of MBD9 under its native promoter (2kb upstream of TSS) was cloned into pDONR221 gateway plasmid through BP recombination (Invitrogen) as previously described ^49^. Aspartic acid 1209 of MBD9 was mutated to Alanine through GAT→GCT DNA sequence change using the New England Biolabs (NEB) Site Directed Mutagenesis Kit and D1209A Mutagenic primers listed in Supplemental Table 1. The D1209A MBD9 sequence was cloned into destination gateway plasmid pMDC99 through LR recombination (Invitrogen) and verified by Sanger Sequencing. D1209A MBD9 in pMDC99 and WT MBD9 in pMDC99 were transformed into *Agrobacterium tumefaciens* GV3101 strain via electroporation. *mbd9-1* plants were transformed with either WT MBD9 or D1029A MBD9-containing plasmids through the floral dip method. Stable transformants were identified through selection on Hygromycin B-containing media (35 μg/mL) and selfed until homozygosity of the transgene. T4 generation plants were used for all described experiments.

### Abscisic Acid (ABA) treatment and Germination Efficiency

For Chip-Seq and RNA-Seq, above-ground tissue of plate-grown seedlings was thoroughly sprayed with an ABA or Mock solution (∼3 mLs solution per 150mm diameter plate). The ABA solution contained 20 µM (+/-) Abscisic Acid (ABA) (Phytotech Labs) with 0.01% Silwet® L-77 (bioWORLD PlantMedia) in water, making the active form (+) ABA 10 µM in final concentration. The Mock solution contained an equal volume of DMSO (ABA solvent) and 0.01% Silwet® L-77 in water. Sprayed seedlings were incubated at 20°C in growth chambers and collected after 4 hours, approximately 9 hours after artificial dawn. For ABA germination experiments, half-strength Murashige and Skoog (MS) media plates were supplemented with ABA to a final concentration of 0.6 µM (+/-) Abscisic Acid (ABA) (Phytotech Labs), making the active form (+) ABA 0.3 µM. Seeds were plated at a low density (∼50-80 per 150mm diameter plate) to ensure full contact with ABA-containing media and stratified and grown as described above. The emergence of root and shoot tissue was tracked on a semi daily basis. Seeds of each genotype were also germinated on standard ½ MS media simultaneously as a control for each iteration of the germination experiment.

### RNA Extraction

Total RNA was isolated from three biological replicates of plate-grown seedling tissue (100 mg per replicate) using the RNeasy plant mini kit (Qiagen). Isolated RNA was treated with Ambion TURBO™ DNase (Invitrogen) using manufacturer’s recommendations.

### Quantitative Real-Time PCR (RT-qPCR)

First strand cDNA was synthesized from DNase-treated RNA with LunaScript® RT SuperMix Kit (NEB). Prepared cDNA was amplified in Real Time utilizing the StepOnePlus real-time PCR system (Applied Biosystems) and PowerUp SYBR Green Master Mix (Applied Biosystems) as a detection reagent. The 2^-ΔΔCt^ method was used to determine differential expression, with the *PP2A* gene (*AT1G13320*) used as the internal control ^59^. Primers against exon junctions for genes *COR15A* (*AT2G42540*) and *ANAC72* (*AT4G27410*) were used and described in Supplemental Table 1. Their expression was compared across Mock and ABA treatments to ensure sufficient treatment with ABA.

### RNA-Seq library generation and sequencing

500 nanograms of DNase-treated RNA from RNA Extraction was used as input for the RNA-Seq library prep. The QuantSeq FWD 3’ mRNA-Seq V2 Library Prep Kit with UDI (Lexogen) was used to make sequencing libraries with manufacturer’s recommendations. Resulting libraries were pooled and sequenced using paired-end 150 nt reads on an Illumina NovaSeq 6000 instrument.

### RNA-Seq data analysis

Read 1 from RNA-seq was trimmed with Trimmomatic using standard single end (SE) parameters ^60^ and then aligned to the Col-PEK genome assembly ^61^ using the STAR alignment software ^62^. Picard MarkDuplicates function with REMOVE_DUPLICATES=T parameter was used to remove sequencing duplicates (https://broadinstitute.github.io/picard/). featureCounts was then used to convert alignment files into count matrix files for downstream analyses ^63^. DESeq2 was used to identify differentially expressed genes fold change of 1.5 or greater (log2FoldChange <= -0.6 or >= 0.6) and a Benjamini-Hochberg (BH) adjusted p-value <0.05 as reported by DESeq2 ^64^ .

### ChIP library generation and sequencing

The enhanced ChIP-Sequencing protocol was followed as previously described in ^65^ with the following modifications: Per replicate, 200 mg of 7-day old seedling tissue (above ground tissue only) was harvested, crosslinked in 1% formaldehyde, and flash frozen in liquid nitrogen. Tissue was ground in a mortar and pestle, homogenized into a slurry in 250 µl of Buffer S and moved directly to sonication. Post sonication, lysate was diluted 10x with Buffer F for use in pulldowns. 50 µl of diluted lysate was saved as input. αH2A.Z (2 μg/mL), αMBD9 (2 μg/mL) or αPIE1 (3 μg/mL) antibodies were added to diluted lysates and incubated overnight at 4°C on a nutator for end-over-end mixing. The H2A.Z antibody used here is described by Deal et al. in 2007 ^66^, and MBD9 and PIE1 antibodies described above. 20 µl of Protein G Dynabeads (Invitrogen) per replicate were washed in excess Buffer F, resuspended to their original volume and added to each pulldown. Samples were incubated with beads at 4°C on a nutator for 2 hours. ChIP DNA was extracted via the MinElute PCR Purification Kit (Qiagen) and sequencing libraries were made using the ThruPLEX DNA-Seq Kit (Takara). Two replicates of each genotype/treatment were collected for sequencing. Resulting libraries were sequenced in two batches: ABA- and Mock-treated libraries were pooled and sequenced using paired-end 150 nt reads on an Illumina NovaSeq 6000 instrument while untreated plants were pooled and sequenced using paired-end 150 nt reads on an Illumina NextSeq 500.

### ChIP sequencing data analysis

Sequencing reads were aligned to the Col-PEK genome assembly ^61^ with Bowtie2 ^67,68^ using --local --very-sensitive --no-mixed --no-discordant --phred33 parameters. Samtools -view bS, -index, and -sort functions were used to process resulting alignment files ^69^. Picard MarkDuplicates function with REMOVE_DUPLICATES=T parameter was used to remove sequencing duplicates (https://broadinstitute.github.io/picard/). deepTools2 bamCoverage function with parameters --normalizeUsing RPKM and --extendReads were used to create Read Per Kilobase Million (RPKM) normalized bigwig files for visualization ^70^. htseq-count with -r pos -m union -s no --nonunique all parameters were used to convert processed BAM files into count matrix files for downstream analyses ^71^. The R package DESeq2 was used to identify genes with significantly changed enrichment of H2A.Z, MBD9 and PIE1 for each genotype or condition comparison mentioned in the text ^64^. After running DESeq2 analysis, results were further filtered to exclude genes with low baseMean (<20 for H2A.Z, <5 for MBD9/PIE1).

### Data visualization

Unless otherwise noted, heatmaps and average plots were generated using the Seqplots software using RPKM normalized bigwig files and BED files containing the genomic coordinates of all or a specified subset of Col-PEK genes ^72,73^). For **Fig. 1c**, deepTools2 computeMatrix and plotHeatmap were used in lieu of Seqplots to generate average plots ^70^. Integrative Genomics Viewer (IGV) was used to visualize specific loci in **Fig. 3h** ^74^. All other plots including volcano plots, scatter plots, bubble plots, box plots, and germination rate plots were made in R utilizing ggplot2 ^75^.

### Publically available data

Raw data for histone post-translational modifications (PTMs) H3K9Ac, H3K14Ac, and H3K27me3 presented in **Fig. 1c** were obtained from the NIH Sequencing Read Archive database under the following accessions, respectively; SRR5011170, SRR5011158 and SRR5278091 ^76^. The two replicates of Global Run-on Sequencing (Gro-Seq) data from **Supp. Figure 6** are available under accessions SRR3647034 and SRR3647034. Raw data were processed as above into RPKM normalized bigwig files for visualization.

### Gene Ontology (GO) Analysis

The Bioconductor package ViSEAGO ^77^ was used for Gene Ontology (GO) Analysis following the general tutorial provided by Brionne, A et al. 2024^78^. Briefly, select gene lists were mapped to their corresponding Entrez ID utilizing the select function from the AnnotationDbi package in R ^79^. ViSEAGO::create_topGOdata was used with ont=“BP”, nodeSize=5 parameters to assign ‘Biological Process’ (BP) to genes. Then runTest from the R package topGO was used with algorithm =“elim”, statistic = “fisher” parameters to determine statistical enrichment of GO terms using a Fisher’s Exact Test (p-value threshold < 0.01) ^80^. Enriched GO categories were condensed based on Semantic Similarity (SS) clustering using distance=“Wang” and aggreg.method=“ward.D2” parameters from the ViSEAGO::GOclusters_heatmap function.

### Statistics

For ChIP-Seq and RNA-Seq data analysis, differentially enriched regions and differentially expressed genes were defined as genes with a fold change of 1.5 or greater (log2FoldChange <= -0.6 or >= 0.6) and a Benjamini-Hochberg (BH) adjusted p-value <0.05 as reported by DESeq2. Two independent biological replicates for ChIP-Seq and three for RNA-Seq were performed independently from extraction to sequencing. Unless otherwise noted, two-sided Welch Two Sample t-tests were performed for all log2 Fold Change comparisons of the resulting data. Germination efficiency values described in **Fig. 4e** represent the average germination of genotypes across two independent germination experiments, where the germination of an average of 68 seeds per plate were counted. A two-sided analysis of variance ANOVA was performed to determine genotype-specific germination rate differences.

### Insoluble Chromatin Isolation and Western Blotting

To isolate insoluble chromatin associated proteins, we followed the isolation protocol described previously ^48^. Briefly, 1 gram of 7-day-old, plate grown seedling shoots were collected, ground in liquid nitrogen, and homogenized in 5 ml of modified Honda buffer without PMSF. Homogenates were filtered through 70 uM cell strainers and centrifuged at 1,500g at 4°C for 20 minutes. Pellets were washed three times in 2 ml modified Honda buffer and once in 2 ml 1x PBS with 1 mM EDTA, with spins at 1,500 for 5 minutes at 4°C. Resulting chromatin fractions were resuspended in equal volumes of 1X Laemmli Buffer. Western blotting was performed as previously described ^49^, using 1:1000 dilutions of αH2A.Z and αPIE1 antibodies and 1:500 of αMBD9 antibodies. ECL was used as a chemiluminescent detection reagent (Thermo Fisher) and scanned for signal using ChemiDoc MP imaging system (BioRad).

### Data availability

ChIP-Seq and is available under the GEO accession GSE269827 and RNA-Seq data is available under the GEO accession GSE269876.

## Supplementary information

**Supp. Table 1.**
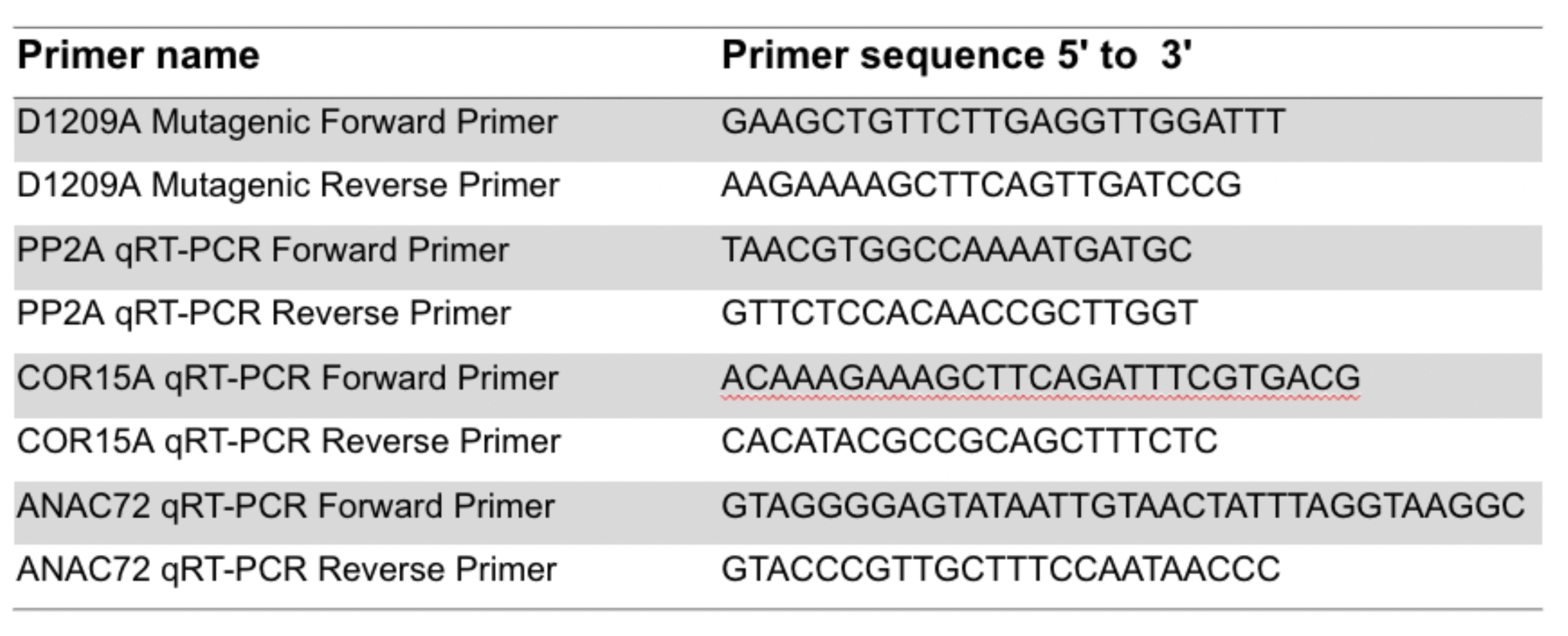
shows primer sequences mentioned in the text.

**Supp. Fig. 1.**
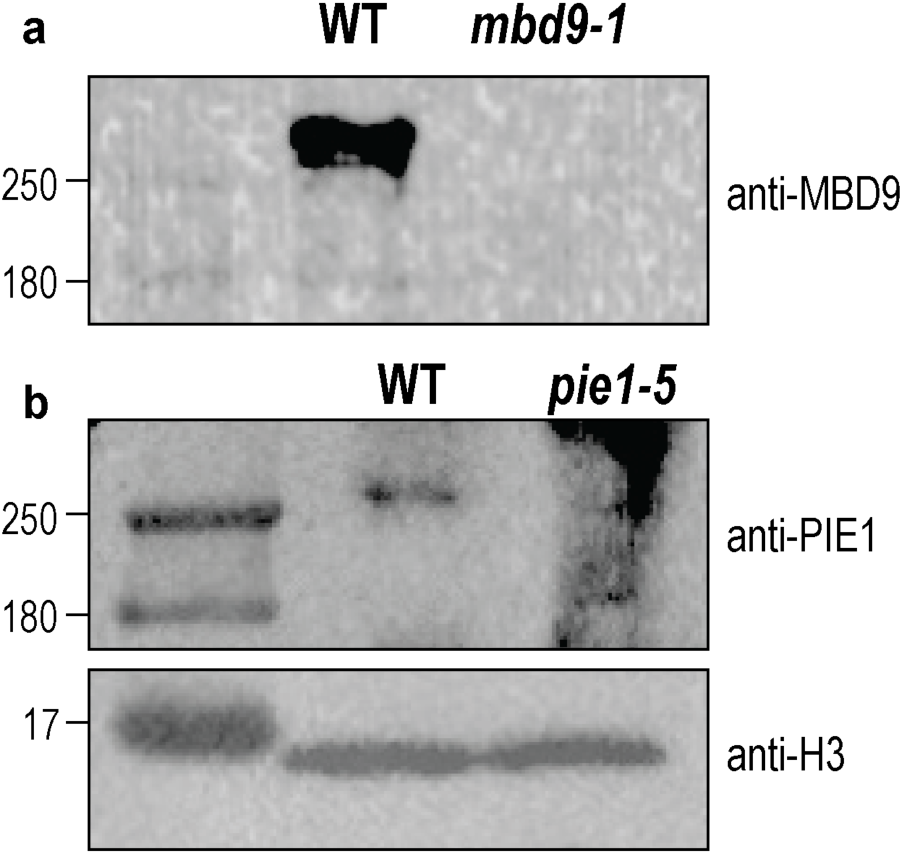
PIE1 and MBD9 antibodies recognize their respective proteins by Western Blot (a) Western blot against MBD9 in WT and mbd9-1 plants. Nuclear proteins were extracted following as described in the methods for insoluble chromatin isolation nuclei isolation and Western Blotting. The band in WT is the expected size for MBD9, which is around 270 kDA, and is not detected in the *mbd9-1* extract. (b) Western blot against PIE1 in WT and *pie1-5* plants. The band in WT is the expected size for PIE1, which is around 250 kDA and is not detected in *pie1-5* extract.

**Supp. Fig. 2.**
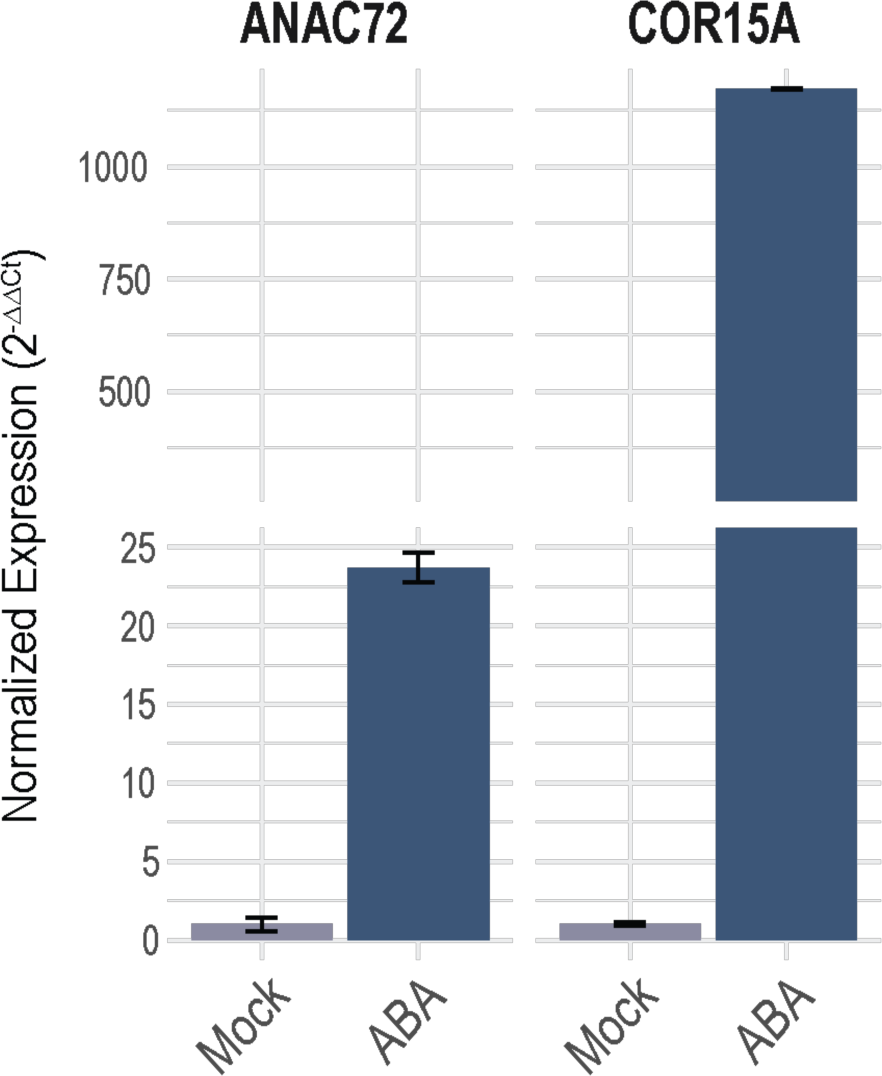
Bar graph illustrating WT qRT-PCR response to 10 uM Abscisic Acid (ABA) at two genes, *COR15A* (*AT2G42540*) and *ANAC72* (*AT4G27410*) compared to Mock treatment. Normalized expression represents the change in expression of these genes compared to both an internal control (*PP2A*;*AT1G13320*) and relative to Mock treatment (no ABA). Clearly, WT plants have a strong response to ABA, with relative transcript level increasing 25x for the *ANAC72* gene and 1000x for the *COR15A* gene.

**Supp. Fig 3.**
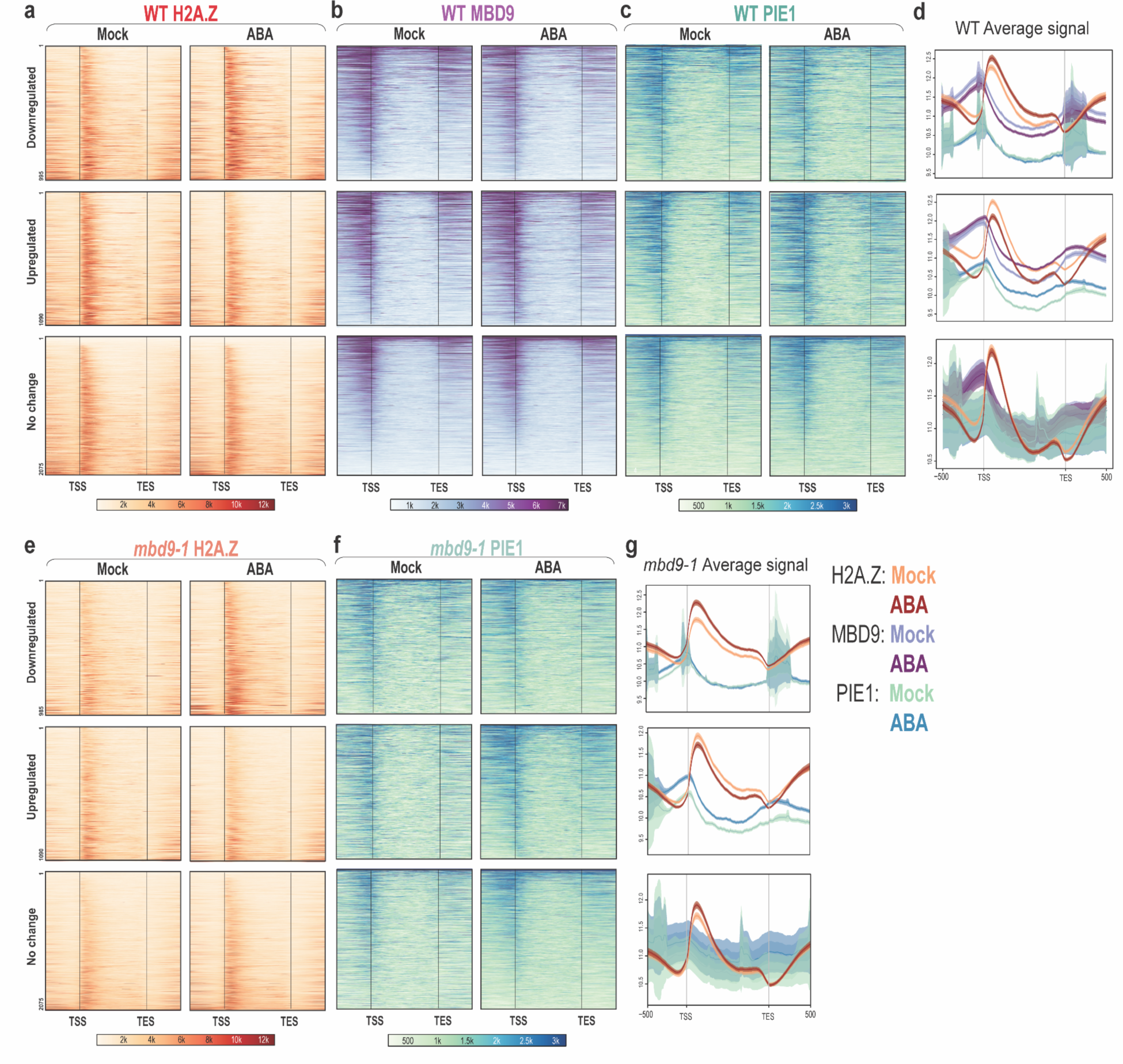
shows chromatin dynamics of WT and *mbd9-1* plants at C2 genes. **(a)** Heatmap showing WT H2A.Z localization at 995 C2 genes that are Downregulated (top), 1090 genes that are Upregulated (middle) and genes that have no change by RNA-Seq (bottom). H2A.Z increases slightly at downregulated genes, decreases at upregulated genes, and does not change at unchanged genes. **(b)** Heatmap with the same groups as in (a) but showing MBD9 localization on genes. There is a decrease of MBD9 at downregulated genes, a slight increase at upregulated genes, and no change at unchanged genes. **(c)** shows the same trends with PIE1 data. All data are RPKM normalized. **(d)** Average plots illustrating the average change in signal at these genes. Profiles are plotted on a log2 scale for easy visualization. **(e)** Heatmaps illustrating H2A.Z changes in *mbd9-1* plants. Note that heatmaps are on the same scale as WT plots in (a-c). mbd9-1 plants still show gains in H2A.Z at the same C2 genes as in (a), and loss of H2A.Z in upregulated genes from (a). **(f)** Heatmaps illustrating PIE1 change in *mbd9-1* plants. The most notable change is at Upregulated genes, where more PIE1 is localized after ABA treatment. **(g)** Average plots illustrating average change in H2A.Z and PIE1 in these three groups of genes in *mbd9-1* plants. Profiles are plotted on a log2 scale for easy visualization.

**Supp Fig. 4.**
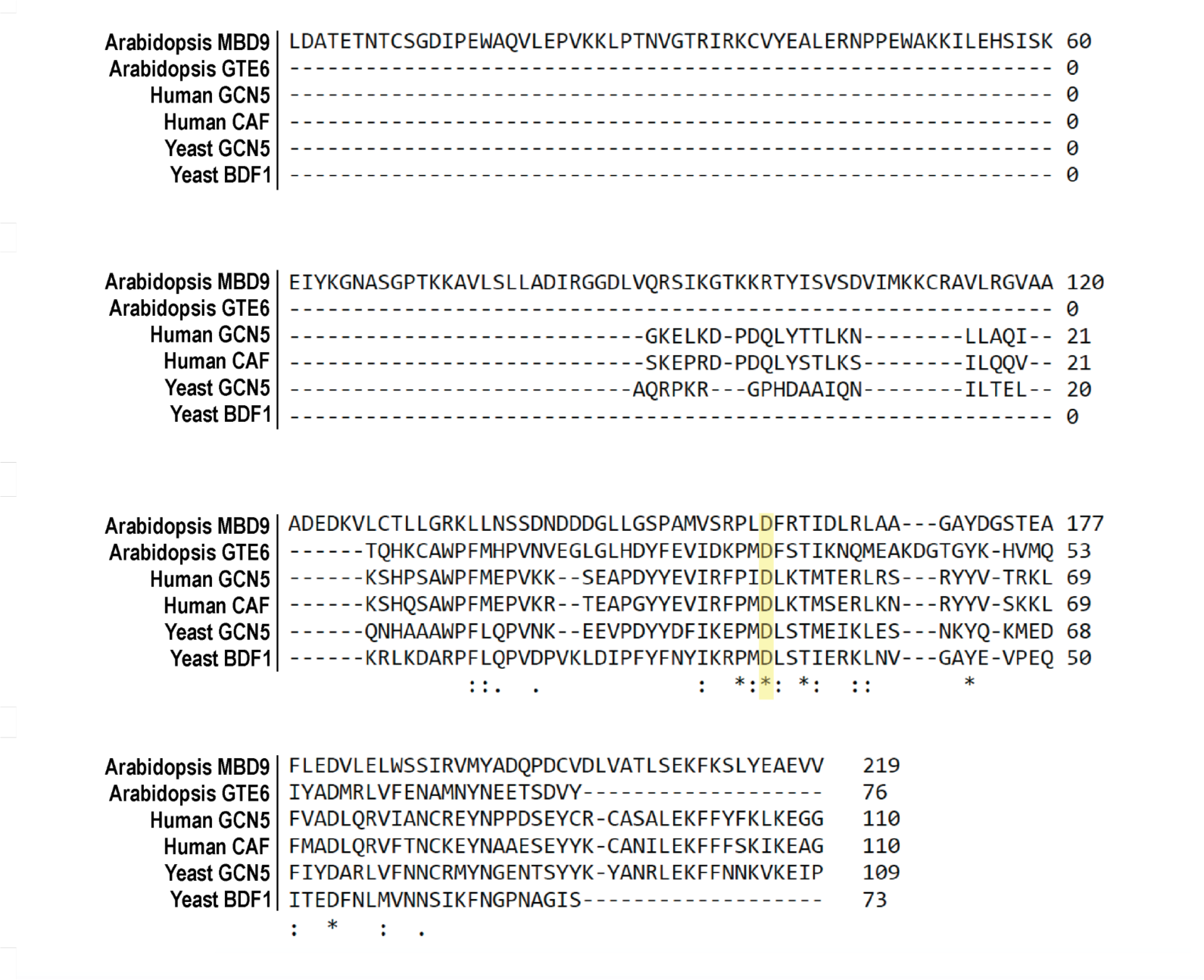
shows ClustalOmega (1.2.4) multiple sequence alignment of the Bromodomains from Arabidopsis *MBD9 (AT3G01460)*, Arabidopsis *GTE6* (*AT3G52280*), Human GCN5 (ENST00000688188.1), Human p300/CBP-associated factor (CAF) (ENSG00000114166), *Saccharomyces cerevisiae* (Yeast) GCN5 (YGR252W), and Yeast BDF1 (YLR399C). ” * ”indicates identically conserved residues, , ” : ” indicates strongly similar residues, ” . ” indicates residues with weakly similar properties, and no symbol indicates residues that are not conserved. The highlighted Aspartic Acid (D) correlates to D1209A in the total *MBD9* sequence. This residue is identically conserved across all compared Bromodomains here.

**Supp. Fig 5.**
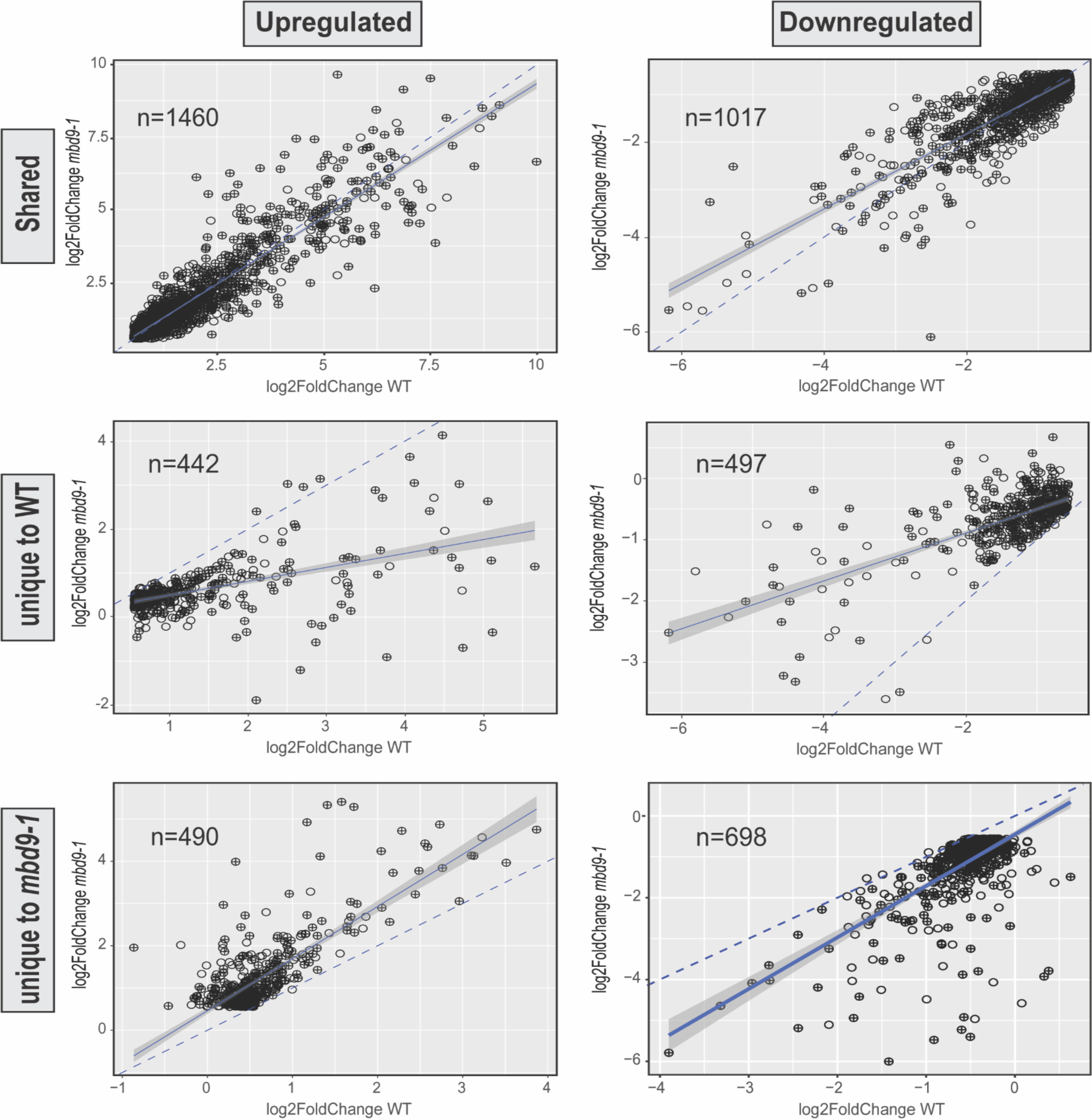
shows scatter plots illustrating correlation of WT and *mbd9-1* RNA-Seq reponses at gene groups from Fig. 5c. Here, dashed lines indicate hypothetical 1:1 relationship between WT and *mbd9-1* responses whereas solid blue lines indicate the trend of the data itself, with gray shadows indicating the standard error of the data. Although there are upregulated and downregulated genes ‘unique’ in both WT and *mbd9-1* plants, many of these points still trend in the same direction but may not be called as statistically significant through DeSeq2 analysis. The group with the most notable deviation from the expected 1:1 ratio is the Upregulated Unique to WT Category (middle left panel) which has some genes that have opposing or little change in *mbd9-1* plants but a significant change in WT plants.

**Supp. Fig 6.**
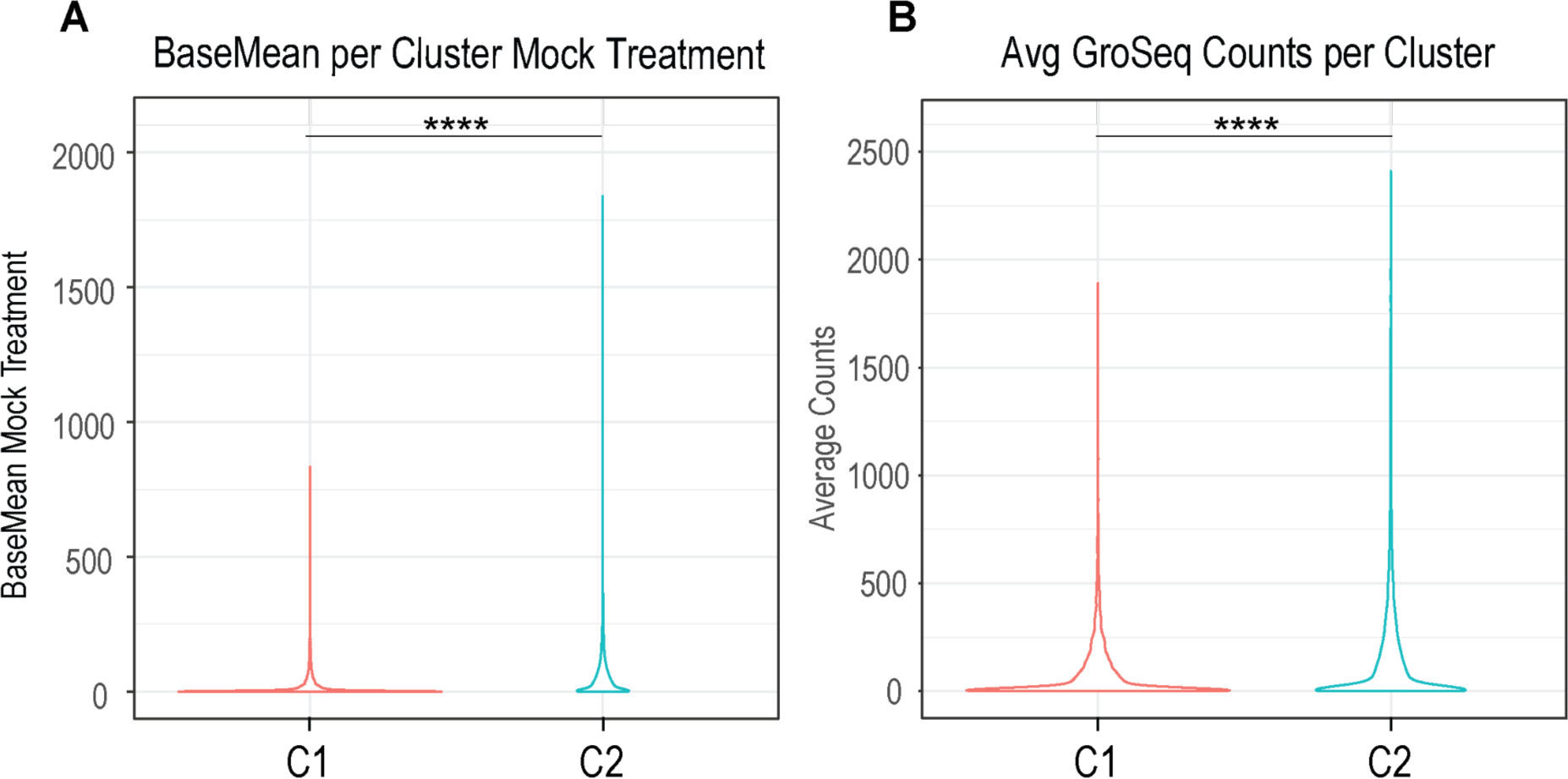
shows quantification of count data from RNA-Seq and publicly available nascent RNA data Global Run-on Sequencing (GRO-Seq) from analogous tissue. **(a)** Violin plot showing differences in RNA-Seq BaseMean (normalized count data outputted from DeSeq2) between Clusters in plants under Mock treatment. C2 has significantly more counts compared to C1, indicating that under Mock treatment (no ABA), C2 genes are more transcribed than C2 genes (unpaired, two-sided Wilcoxon rank-sum test,****= p-value<0.001). **(b)** Violin plot showing average differences in Gro-Seq counts from FeatureCounts across two replicates between Clusters in untreated, 6-day-old WT Arabidopsis seedlings. Again, C2 has significantly more counts compared to C1, indicating that under normal conditions, C2 genes are more transcribed than C2 genes (unpaired, two-sided Wilcoxon rank-sum test,****= p-value<0.001).

## Notes

### Competing Interest Statement

The authors have declared no competing interest.

### Summary of Updates

Reference DOI added to a co-submitted manuscript on bioRxiv.

